# Interactions of nuclear transport factors and surface-conjugated FG nucleoporins: Insights and limitations

**DOI:** 10.1101/435651

**Authors:** Ryo Hayama, Mirco Sorci, John J. Keating, Lee M. Hecht, Joel L. Plawsky, Georges Belfort, Brian T. Chait, Michael P. Rout

## Abstract

Protein-protein interactions are central to biological processes and the methods to thoroughly characterize them are of great interest. *In vitro* methods to examine protein-protein interactions are generally categorized into two classes: in-solution and surface-based methods. Here, using the multivalent interactions involved in nucleocytoplasmic transport as a model system, we examined the utility of three surface-based methods in characterizing rapid interactions involving intrinsically disordered proteins: atomic force microscopy, quartz crystal microbalance with dissipation, and surface plasmon resonance. Although results were comparable to those of previous reports, the apparent effect of mass transport limitations was demonstrated. Additional experiments with a loss-of-interaction mutant variant demonstrated the existence of additional physical phenomena and an uncharacterized binding mode. These results indicate the binding events that take place on the surface can be quite complex, suggesting particular care must be exercised in interpretation of such data.

## Introduction

Protein-protein interactions are at the core of any biological system and regulate essential cellular functions; measuring their characteristics, such as stoichiometry, affinity and kinetics, is crucial for understanding their biological roles. There are multiple *in vitro* methods to characterize protein-protein interactions, most popular of which are surface-based such as enzyme-linked immunosorbent assay (ELISA) and surface plasmon resonance (SPR). These surface-based methods have been applied to a wide range of protein-protein interactions: from well-defined antigen-antibody interactions to those involving intrinsically disordered proteins (IDPs), a major class of proteins involved in various functions, many of whose detailed behaviors are still being characterized [1, 2]. Here, we examined in detail the applicability of select surface-based techniques to a complex system involving IDPs, specifically the one involved in the nucleocytoplasmic transport mediated by nuclear pore complexes (NPCs) [3–6].

NPCs are the sole conduits across the nuclear envelope; macromolecular exchange between the nucleoplasm and the cytoplasm occurs in their central tube, which is lined with extensive regions of intrinsically disordered FG nucleoporins (FG Nups), so-called because of the presence of multiple phenylalanine-glycine (FG) repeat motifs. It is generally agreed that protein-protein interactions between the FG repeat motifs in FG Nups and transport factors (TFs) are central to selective and rapid nuclear transport across the NPC [4, 7]; however, the exact physical mechanism of nuclear transport remains poorly characterized. There have been many reports on measurements of the strengths and modes of interactions between FG Nups and TFs [8–19]. The methods employed to study the FG-TF interaction vary, although most of them utilize surface-based systems, including microtiter plate and beads binding assays [8–12, 20], atomic force microscopy (AFM) [21–23], bio-layer interferometry [24], SPR [14–16], and quartz crystal microbalance with dissipation (QCM-D) [13, 25–27]. Many of these methods report low micromolar to nanomolar dissociation constants (*K_D_*s) for the binding affinity between FG Nups and TFs [13–16, 20]. Although FG Nups are grafted onto a tubular surface in the actual NPC, and so such surface grafted systems seem consistent with the situation *in vivo*, the strong affinities observed in these experiments are at odds with the fast transport rates (∼5-20 ms translocation time) seen *in vivo* [28–34]. Recently, we and others have reported in-solution affinities between TFs and individual FG motifs, whose per-FG-motif *K_D_*s were in the millimolar range, compatible with the rapid kinetics of TF translocation through the NPC [17–19]. We also found that multiple low-affinity interactions can yield a higher overall interaction specificity than monovalent ones without compromising a high on-off rate of individual FG motifs [19]. Thus, one motivation of this work was to investigate the cause of these discrepancies in affinity measurements conducted by various methods.

As outlined by Schuck and Zhao, analysis of multivalent interactions by surface-based methods require extra care because of potential complexities of the binding mechanism on the surface; in some cases rendering the results “impossible to realistically interpret” [35]. In addition, (i) it is often difficult to quantify the amount or the density of protein conjugated to the surface; (ii) conjugated surfaces usually have an inhomogeneous distributions of ligands [35]; (iii) the degree and the effect of analyte retention on the surface after a binding experiment is often not assessed; (iv) mass transport limitations can significantly affect measurements of binding kinetics when using SPR and QCM-D, and are often overlooked [35–37]; (v) change in protein conformation or denaturation upon binding to surfaces can occur [38, 39], and (vi) macroscopic effects of surface crowding, especially in the context of multivalency, are not trivial to adequately address and quantify for surface-based methods [35].

A well-characterized model FG-TF system, the FG repeat region of Nsp1 (an FG Nup containing at least 35 repeats), Nsp1FG, and Kap95 (a TF), was used to investigate some of the complexities outlined above. Nsp1 is the most abundant FG Nup in budding yeast (*S. cerevisiae*) and consists of two distinct FG regions: an N-terminal FG-type repeat region rich in polar amino acids such as asparagine, and the central FxFG-type repeat region that is rich in charged amino acids and has a highly conserved repeat sequence [40]. Kap95 belongs to a major class of TFs called karyopherins, which mediates the main canonical nuclear import pathway. Collectively, Kap95-Nsp1 interactions mediate a large fraction of nucleocytoplasmic trafficking [41]. Karyopherins are comprised almost entirely of superhelical alpha-solenoids. Co-crystal structures with FG repeat peptides indicate that their interaction is mediated through the insertion of the phenylalanine from an individual FG motif into a cleft formed between adjacent alpha-helices on the outer surface of the solenoid [10, 42–44]. Kap95 is thought to have multiple binding sites for FG motifs, though the exact number of binding sites is unknown (its ortholog, CRM1, is reported to have eight FG-interaction sites [45]). Interactions between Nsp1 and Kap95 have been extensively characterized by various methods, including crystallography and NMR [10, 12, 15–17, 20, 42–44, 46, 47]. In this study, the Nsp1-Kap95 interaction was analyzed by three commonly used surface-based methods: AFM, QCM-D and SPR. AFM has been employed to characterize the nanomechanical properties of proteins, including FG Nups [21–23]. QCM-D and SPR can monitor binding of analytes to ligands anchored to the surface of their sensors in real time and have been used to study FG Nups [13–16, 25–27]. The results obtained in this study qualitatively replicated earlier results using the same methods [13–16, 21–23, 25–27]; however, our results implicate macroscopic and likely non-biological physical events, notably mass transport limitations, which must therefore be accounted for in any analysis utilizing similar methods.

## Materials and Methods

### Materials

HS-(CH_2_)_11_-EG_3_-OH (hPEG) and HS-(CH_2_)_11_-EG_3_-OCH_3_ (mPEG) were purchased from Prochimia (Sopot, Poland). (3-aminopropyl)-trymethoxysilane was purchased from Gelest, Inc. (Morrisville, PA). Solution P: 18 mg/ml phenylmethylsulfonyl fluoride (PMSF), 0.3 mg/ml pepstatin A in ethanol, cOmplete™ EDTA-free protease inhibitor cocktail (PIC) (Roche Applied Science). All other chemicals were purchased from Sigma-Aldrich Co. (Saint Louis, MO), as ethylenediaminetetraacetic acid (EDTA), sodium dodecyl sulfate (SDS), 2-mercaptoethanol (βME), tris-(2-carboxyethyl)phosphine (TCEP). The buffer solutions prepared for the experimental work were: (i) PB: 0.01 M phosphate buffer, pH 7.4; (ii) PB-E: PB and 0.01 M EDTA; (iii) PBS: PB and 2.7 mM KCl and 137 mM NaCl_2_, pH 7.4; (iv) PBS-ET: PBS, 10 mM EDTA and 5 mM TCEP; (v) TB: 20 mM HEPES, 110 mM KOAc, 2 mM MgCl_2_, 10 μM CaCl_2_ and 10 μM ZnCl_2_, pH 7.4; (vi) TBT: TB and 0.1% v/v Tween® 20; (vii) TBT-D: TBT and 5 mM DTT; (viii) TBT-PVP: TBT and 0.3% (w/v) polyvinylpyrrolidone; (ix) HUT: 50 mM HEPES, 8 M urea and 0.5% Tween® 20. All buffers were prepared fresh and filtered through a 0.22 μm polyethersulfone membrane Express™ Plus from Millipore (Billerica, MA). Compressed N_2_, He and O_2_ were supplied by Airgas (Albany, NY).

### Protein purification

The expression and purification of Nsp1FG construct containing C-terminal cysteine residue followed by hexa-histidine tag was conducted as previously described [19, 20]. Cells expressing FG constructs were thawed and resuspended in 30 ml of lysis buffer (20 mM HEPES-KOH pH 7.4, 150 mM NaCl, 1% solution P, 1x PIC) including 8 M urea. Cells were lysed by Microfluidizer® (Microfluidics) and the lysate was clarified by centrifugation for 1 h at 192,838 g in a Type 50.2 Ti rotor (Beckman) at 4°C. The supernatant was filtered through a 0.22 μm or 0.45 µm filter depending on its viscosity and was loaded into a 10 ml TALON Sepharose resin (GE Healthcare) column. The resin was washed with 50 ml of (i) lysis buffer containing 8 M urea, (ii) lysis buffer containing 3 M urea, and with (iii) lysis buffer containing 10 mM imidazole and 3 M urea. The protein was eluted in lysis buffer containing 250 mM imidazole and 3 M urea. Eluted fractions were analyzed by SDS-PAGE. Fractions containing the protein were pooled and concentrated by centrifugal concentrators with 3 kDa molecular weight cutoff (EMD Millipore). The sample was polished by gel filtration (120 ml Superdex200) in 20 mM HEPES-KOH pH 7.4, 150 mM NaCl, 1 mM EDTA, 50 μM TCEP, 10% glycerol. Fractions containing the protein were pooled, concentrated, and, dialyzed against 20 mM HEPES-KOH pH 7.4, 300 mM NaCl, 1 mM EDTA, 50 μM TCEP, 20% glycerol. The sample was frozen with liquid nitrogen and stored at −80°C.

Cells expressing GST-Kap95 were thawed and resuspended in TBT-D. The cells were lysed, and the lysate was cleared in the same manner as FG constructs above. Cleared lysate was loaded onto a glutathione-sepharose (GE Healthcare) column (GST-column) equilibrated with TBT-D. The resin was washed with 100 ml of (i) TBT-D, (ii) TBT-D containing 500 mM NaCl, and (iii) TBT-D. GST-Kap95 was eluted with 90 ml of 100 mM Tris-HCl pH 8.5, 20 mM glutathione. Eluted fractions were analyzed by SDS-PAGE and fractions containing GST-Kap95 were pooled. The pooled sample was then dialyzed against Buffer A (20 mM HEPES-KOH pH6.8, 150 mM KCl, 2 mM MgCl_2_). GST was cleaved off by biotinylated thrombin (EMD Millipore), and enzyme was removed by streptavidin sepharose resin (EMD Millipore) following the manufacturer’s instructions. Cleaved GST was removed by the regenerated GST column, and the flow through containing Kap95 was collected. The sample was then concentrated by 30k MWCO Centricon (Millipore) and gel filtered in Buffer A containing 10% glycerol. Fractions containing Kap95 were pooled, concentrated by 50k MWCO Centricon (Millipore) and dialyzed against Buffer A supplemented with 5 mM DTT and 20% glycerol overnight. The sample was frozen with liquid nitrogen and stored at −80°C.

### Atomic force microscopy (AFM)

All force measurements were performed using the MFP-3DTM atomic force microscope (Asylum Research, Santa Barbara, CA) and the collected data were analyzed using IGOR Pro 6 (WaveMetrics, Inc., Lake Oswego, OR). Gold-coated surfaces were prepared by coating glass coverslips (0.20 mm, Corning, NY) with 15 nm of titanium (Ti, 99.999% International Advanced Materials, Spring Valley, NY) followed by 50 nm of gold (99.999%, International Advanced Materials) using the electron beam evaporator under a pressure of less than 10-6 Torr. Nsp1FG solutions were first centrifuged for 10 min at 90,000 rpm in a TLA 100 rotor (Beckman) at 4°C, to remove possible precipitates occurred during protein storage. Two different protocols were followed in order to form either a sparse or a dense layer of Nsp1FG on the surface, sNsp1FG or dNsp1FG, respectively. For sparse sNsp1FG samples, gold-coated surfaces were: (i) Passivated with 2 mM hPEG for 5 min at RT; (ii) rinsed with EtOH and dried with N_2_; and finally (iii) incubated with reduced Nsp1FG solution for 48 h at 4°C. The reduced Nsp1FG solution was prepared by diluting the centrifuged Nsp1 sample to a concentration of 10 μg/ml in PBS and incubated with 1 mM TCEP for 1 h at 4°C. For dNsp1FG samples, we modified a previously published protocol [48]. Gold-coated surfaces were: (i) incubated with reduced Nsp1FG solution for 1 h at 4°C; (ii) soaked in 8 M urea in TBT for 30 min at RT; and finally (iii) incubated with 2 mM hPEG, 5 mM TCEP and 8 M urea for 48 h at 4°C. The reduced Nsp1FG solution was prepared by buffer exchanging the centrifuged Nsp1FG sample into TBT using Microcon centrifugal filter units (Millipore Corporation, Bedford, MA), concentrated to ∼0.4 mg/ml and incubated with 5 mM TCEP for 1 h at 4°C.

For sNsp1FG, single molecule force spectroscopy (SMFS) scans [49–53] were obtained in the so-called “Force–Volume (FV) mode” [54, 55] using biolevers RC-150-VB (Olympus, Center Valley, PA). The cantilevers were calibrated before each experiment, using previously described 3-steps procedure [56]. Spring constants were within 10% error from the ones supplied by the manufacturer. A FV-data set consisted of an array of 1,600 (40 x 40) force measurements, scanning in contact mode at 1 μm/s an area of 5 x 5 μm^2^, with each pixel point spanning an approximate width of 125 nm in both X and Y. The trigger force was set to 500 pN.

For dNsp1FG, SMFS scans were obtained in FV mode using gold coated silicon nitride cantilevers carrying a borosilicate glass sphere, diameter = 10 μm, and a nominal spring constant of 60 pN/nm (Novascan, Ames, IA). The cantilevers were calibrated before each experiment. A FV-data set consisted of an array of 900 (30 x 30) force measurements, scanning in contact mode at 1 μm/s an area of 5 x 5 μm^2^, with each pixel point spanning an approximate width of 167 nm in both X and Y. The trigger force was set to 1 nN.

### Quartz crystal microbalance with dissipation (QCM-D)

Frequency and dissipation measurements were performed using an E4 Auto system and its standard flow module QFM 401 (Biolin Scientific/Q-Sense, Linthicum, MD). All data were analyzed using the Sauerbrey model [57] for rigid layer coupled to the QCM-D sensor, or the Voigt-Voinova model [58] when the layer was characterized by a viscoelastic behavior. The latter is a build-in tool in QTools software (Biolin Scientific/Q-Sense, Linthicum, MD), and the fitting parameters are (i) the thickness, (ii) the elastic shear modulus and (iii) the shear viscosity of the layer.

Polished silicon dioxide quartz crystals with fundamental frequencies of 5 MHz (QSX 303, Biolin Scientific/Q-Sense, Linthicum, MD) were washed with acetone for 10 min, isopropanol for 10 min, rinsed with ethanol and dried with N_2_. Organic contaminants were removed by plasma cleaning (Atomflo 400L2 Plasma System, Surfx Technologies, Culver City, CA) at 120 W (30 L/min He, 0.2 L/min O_2_) for 4 min. Plasma treatment also provided a high density of hydroxyl functionalities suitable for subsequent silane modification. Silanization was performed ex-situ by (i) immersing the sensors in 2% v/v (3-aminopropyl)-trimethoxysilane solution in acetone, for 10 min at RT, to introduce reactive amine moieties; (ii) washing with acetone; (iii) rinsing with deionized H2O; (iv) drying with N_2_; and (v) curing at 100°C for 30 min. The sensor was then placed in the QCM-D flow module chamber for (i) succinimidyl 4-(p-maleimidophenyl)-butyrate (SMPB) conjugation, a bifunctional crosslinker containing NHS ester and maleimide moieties, converting the amine-reactive surface into a maleimide-reactive surface; and (ii) Nsp1FG conjugation via a thiol-maleimide coupling. Both reactions were performed in-situ and monitored in real time at 23°C. SMPB conjugation was performed at 100 μl/min by (i) equilibrating the chamber with PB-E for 20 min; (ii) flowing 2 mM sulfo-SMPB in PB-E for 15 min; and (iii) washing with PB-E for 15 min. Sulfo-SMPB formed a rigid (i.e. dissipation ∼0) layer coupled to the surface, hence, according to the Sauerbrey equation, the adsorbed mass is proportional to a normalized decrease in frequency, ΔF/n [Hz]. From the several experiments, the coupling of sulfo-SMBP resulted in ΔF/n ∼-11 Hz, corresponding to a density of ∼2.5 molecules/nm^2^. Nsp1FG coupling was performed at 50 μl/min by (i) equilibrating the chamber with PBS-ET for 20 min; (ii) flowing (and recycling) 0.1-0.2 mg/ml Nsp1FG in PBS-ET for 60 min; and (iii) washing with PBS-ET for 15 min, HUT for 5 min and finally PBS-ET for 15 min. Two passivation steps were finally included to remove any unreacted maleimide group by (i) flowing 5 mM mPEG in PBS-ET for 30 min; (ii) washing with PBS-ET for 10 min; (iii) flowing 50 mM βME in PBS-ET for 30 min; and (iv) washing with PBS-ET for 10 min, HUT for 5 min and finally PBS-ET for 15 min.

Binding/unbinding experiments using Kap95, GST or BSA were performed at 50 μl/min by (i) equilibrating the chamber with TBT-DG for 15 min; (ii) flowing the protein solution in TBT-DG for 10 min (binding step); (iii) washing with TBT-DG for 30 min (unbinding step); and finally (iv) washing with HUT for 5 min and TBT-DG for 15 min (regeneration step). A second set of experiments was performed to measure the affinity between Nsp1FG and Kap95. In order to minimize the effects of mass transfer (e.g. dilution effect when introducing the protein solution in the QCM-D chamber equilibrated with buffer), the runs were performed at 300 μl/min. The binding step was 1 min, while the unbinding step was 15 min.

### Surface Plasmon Resonance (SPR)

All SPR experiments were conducted on an XPR36 system (BioRad) using GLC sensor chips. The sensor was cleaned with 0.5% SDS, 50 mM NaOH, and 100 mM HCl as instructed by the manufacturer. A GLC sensor was activated to create amine-reactive surface using 50 mM sNHS/EDC (sulfo-N-hydroxysulfosuccinimide/1-ethyl-3-(3-dimethylaminopropyl) carbodiimide hydrochloride) chemistry at the flow rate of 30 µl/min for 5 min. 5 mM aminopropyl-maleimide in HEPES Main Buffer (20 mM HEPES-KOH, pH 7.4, 150 mM NaCl, 1 mM EDTA) was applied to the surface to expose maleimides on the surface. C-terminally cysteine-tagged FG constructs were first diluted in HEPES Main Buffer with 20% glycerol, and then they were further diluted in Protein Conjugation Buffer (20 mM PIPES, pH 6.8, 150 mM NaCl, 1 mM EDTA, 0.8 M urea) to produce final ligand (FG Nup) solutions whose concentrations ranged between 8.75 ∼ 140 µg/ml. This two steps dilution ensured that all the final solutions were at the same pH and contained the same percentages of glycerol. FG constructs were then conjugated to the surface by the reaction between the maleimide and the sulfhydryl group on the cysteine residues at 30 µl/min. The reaction was monitored and was manually terminated once the surface conjugation reached a desired level. In the ‘bare surface’ condition, protein was omitted, and plain Protein Conjugation Buffer was run over the surface. The surface was then flushed with 100 mM beta-mercaptoethanol in Protein Conjugation Buffer at 30 µl/min for 8 min to quench unreacted maleimides and to passivate the surface. The conjugation step was completed by equilibrating the surface with HEPES Main Buffer at 30 µl/min for 5 min, after which the instrument was switched to the analyte binding mode (i.e. rotation of the flow channel lid). Prior to the analyte binding experiments, the surface was equilibrated with TBT-PVP at 30 µl/min for 3 min, washed with HUT solution at 30 µl/min for 3 min, and re-equilibrated with TBT-PVP at 30 µl/min for 3 min.

Analytes were diluted in two steps as with the case for FG proteins. They were first with TBT supplemented with 20% glycerol, and then with TBT-PVP to designated concentrations. The concentration of analytes used for the binding experiments are noted in the respective figures. Binding experiment was conducted at 100 µl/min. The duration of association phase was varied for different experiments, ranging between 15 and 120 s. Dissociation was monitored for 10 min. The surface was regenerated after each round of a binding experiment with HUT solution at 30 µl/min for 2 min. The surface was then re-equilibrated with TBT-PVP at 30 µl/min for 3 min before the next round of binding experiment. All binding experiments were conducted at 25°C.

The binding curves were analyzed by XPR36 software. The details of the used models were described previously [59].

## Results

### Establishing robust surface conjugation

In order to study FG Nup behavior and its interaction with TFs and non-specific proteins, we established a robust protocol to immobilize FG Nups on a surface. QCM-D experiments for FG Nups have been conducted previously, where his-tagged FG Nups were conjugated non-covalently to a supported lipid bilayer (SLB) [13, 25–27]. Here, we used alternative protocols for direct, covalent attachment of FG Nups to surfaces because: (i) his-capturing moieties can be mobile when attached to lipids in the SLB, (ii) surface conjugation via his-tag is known to drift (i.e. his-tagged ligands detach and re-attached or drift away due to low affinity non-covalent bonds) [60], and (iii) it is not clear how indirect attachment of FG molecules via SLB would affect the overall surface behavior. Therefore, in this study, a single cysteine residue was inserted at the C-terminal end of the Nsp1 FG domain construct (Nsp1FG) to allow direct, covalent conjugation via the thiol moiety (-SH) to both gold and to chemically activated silica surfaces, so that only the FG molecules covalently attached to a surface can be examined. Following the convention of the SPR literature, the material conjugated onto the surface is designated “ligand” and the material flowing over the surface “analyte”.

The procedure for Nsp1FG conjugation to an amino-activated silica sensor is outlined in Fig 1A and representative QCM-D data is shown in Fig 1B. For gold surfaces, direct conjugation via the C-terminal cysteine was used. Although the analyte binding pattern on gold was similar to that on silica (S1 Fig), we chose silica sensors for protein binding experiments because they exhibited reduced non-specific binding compared to gold sensors (see below).

**Figure 1.**
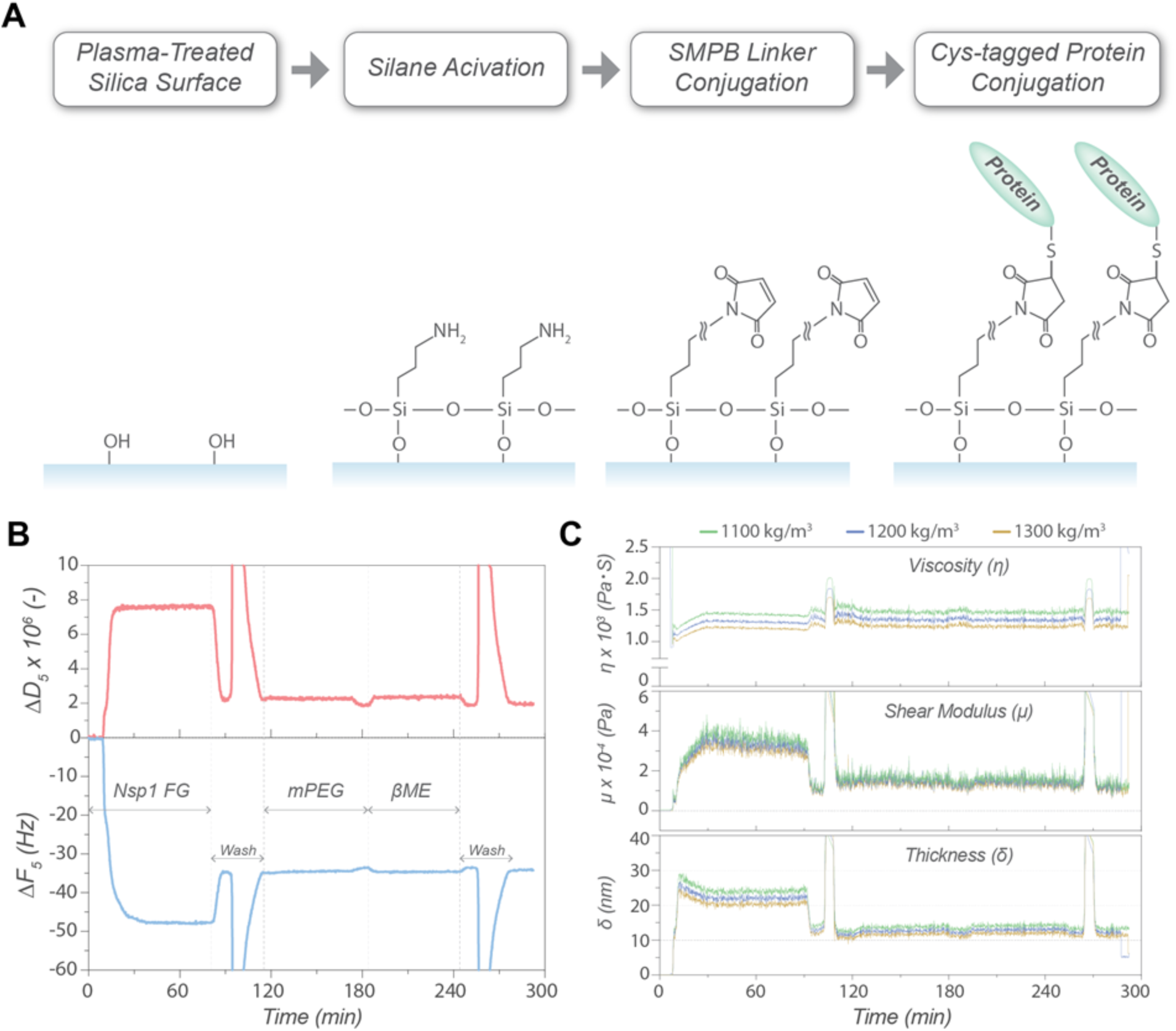
Conjugation of Nsp1FG to a QCM-sensor. (A) Chemical conjugation of Nsp1FG to a silica surface. (B) Representative sensorgram of Nsp1FG conjugation. Changes in resonance frequency (*ΔF_5_*) (bottom, blue) and energy dissipation (*ΔD_5_*) (top, red) for the 5^th^ overtone recorded in real time. Nsp1FG conjugation step, wash, passivation with mPEG, and beta-mercaptoethanol treatment are indicated. (C) Viscoelastic modeling of the Nsp1FG surface at different density estimates (top: viscosity, middle: shear modulus, bottom: layer thickness).

Polyethylene glycol-thiol (PEG-SH) was applied as a passivator [61], followed by a beta-mercaptoethanol treatment to deactivate unreacted maleimide moieties and to minimize non-specific binding. To find an appropriate passivator for protein binding experiments, different PEGs (i.e. number of PEG repeats and chemical groups) were tested against various analytes. Based on the results, a minimum of six PEG repeats were needed to produce an inert surface, while the terminal group did not have a visible effect on the inertness (S2 Fig). Thus, mPEG_6_-C_2_-SH was chosen as the passivator for our experiments. In addition, wash steps to remove any non-covalently bound material from the surface were included (S3 Fig).

To understand the physical properties of a densely packed Nsp1FG layer on a surface, the changes in frequency (*ΔF*) and energy dissipation (*ΔD*) resulting from the Nsp1FG conjugation were subjected to viscoelastic modeling. The Nsp1FG layer was viscoelastic, as indicated by a *ΔD*/*ΔF* ratio larger than 1·10-8 Hz^−1^ [62] and by the spreading of the different harmonics (S4 Fig), consistent with results reported by others [13, 25–27]. The Voigt-Voinova model [58] was used to evaluate changes in layer thickness, shear modulus, and shear viscosity with time (Fig 1C). Varying the layer density estimate from 1,100 to 1,300 kg/m^3^ (i.e. the effective density of the protein layer has to lie between 1,000 kg/m^3^, the value for water, and 1,330 kg/m^3^, the value for proteins, considering a partial specific volume of proteins close to 0.75 ml/g [63–65]) did not significantly affect the estimated parameters. At low Nsp1FG density, the shear viscosity of the layer was close to that of pure water (∼0.001 Pa·s) and increased to 0.00135 ± 0.00003 Pa·s as the Nsp1FG binding progressed when the layer density was set at 1,200 kg/m^3^. The shear modulus equilibrated at 0.012 ± 0.002 MPa and the layer thickness at 12.2 ± 0.5 nm. Although the thickness of the FG layer estimated here was smaller than those previously reported [13, 25–27], this could arise from how the FG-surface was set up, particularly the indirect attachment to the surface via a single lipid bilayer and the non-covalent attachment of FG proteins in those studies. Based on the mass conjugated on the surface (∼1,500 ng/cm^2^, or 25.6 pmol/cm^2^), the Nsp1FG layer behaves more like a polymer brush of 30 kDa PEG, which has shear viscosity below 0.0014 Pa·s and shear modulus below 0.1 MPa, than an agarose hydrogel, which has shear viscosity above 0.0028 Pa·s and shear modulus above 0.2 MPa [66]. Therefore, we were able to produce an FG-surface layer via direct, covalent attachment, whose properties are typical of a viscoelastic brush layer.

### Probing the FG Nup conjugated surface by AFM

To further assess the physical properties of Nsp1FG on a surface, AFM was employed in force mode to interrogate both single Nsp1FG chains (sNsp1FG) grafted at low density, as well as dense layers of Nsp1FG (dNsp1FG). Gold surfaces were chosen for the AFM work because (i) QCM-D experiments showed that the analyte binding properties of the gold surface was similar to that of silica (S1 Fig); and (ii) the absence of a linker between the surface and the protein was preferred when stretching single protein molecules.

Substrate passivation, using hPEG_6_-C_11_-SH, was necessary (i) to minimize non-specific interaction between the gold-coated AFM cantilever tip and the gold surface and (ii) to vary Nsp1FG density on the surface for single molecule stretching. This step was optimized by testing different incubation times of hPEG_6_-C_11_-SH with the gold substrate (S5 Fig): a 5 min passivation step was included in the preparation of the samples discussed below.

All experiments were conducted either in PBS or TBT buffers and the schematic diagram for the AFM experiments is shown in Fig 2A. The Force-Volume (FV) map of sNsp1FG is shown in S6 Fig, where as many as 40 x 40 = 1,600 force curves per sample were collected. Up to a certain extension, ∼50%, stretching of Nsp1FG occurred with forces less than 10 pN, indicating that Nsp1FG is an elastic molecule that can extend itself by ∼50% without any significant force (Fig 2B, normalized data). Peaks showing reproducible stretching events were fitted using the worm-like chain (WLC) model [67, 68]:

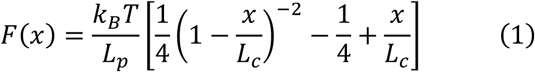

where *F(x)* is the distance-dependent force [kg·m·s^−2^], *x* is the extension length [m], *k_B_* is Boltzmann’s constant [1.3806503·10^−23^ m^2^·kg·s^−2^·K^−1^], *T* is the temperature [K], *L_p_* is the persistence length [m] and *L_c_* is the contour length [m] of the chain (see Fig 2A). Examples of fitted data in PBS are shown in S7 Fig and the fitted values for *L_p_* and *L_c_*, as a function of adhesion force (*F_ad_*), are summarized in Fig 2C. The *L_c_* represents the theoretical length of the stretched molecule. The wide variability of this parameter was due to the random interaction of the AFM cantilever tip with the covalently immobilized sNsp1FG, resulting in partial or full (i.e. ∼250 nm, based on the AA sequence) stretching of the molecule. On the other hand, *L_p_* is a measure of the rigidity of the polymer chain and it is an intrinsic property of the molecule (i.e. independent from *L_c_* or *F_ad_*). The results showed a mean *L_p_* of 1.05 ± 0.29 nm in PBS and 1.10 ± 0.29 nm in TBT buffer (S8 Fig), confirming the elastic behavior upon stretching of sNsp1FG, as previously shown for other FG Nups [21, 22]. Thus, the choice of buffer did not have a significant effect on the measurements. Similar results were obtained when stretching sNsp1FG at different pulling rates in PBS: 1.05 ± 0.29 nm (1 μm/s), 0.80 ± 0.26 nm (2 μm/s), and 0.82 ± 0.22 nm (3 μm/s), confirming that *L_p_* represents the intrinsic flexibility of Nsp1FG and is not caused by the mechanical stretching rate of the molecule (S9 Fig). When normalized to *L_c_*, the retraction force curves collapsed into a “master curve”, confirming homogeneity of the Nsp1FG population and the elasticity of the individual Nsp1FG molecules (Fig 2B). Moreover the *F_ad_* did not exceed 0.2 nN, well below the rupture force of a thiol-Au bond of ∼1.4 nN [69], allowing stretching of Nsp1FG molecule without detachment. The persistence length measured here was larger than that measured for the human Nup153 FG region [21, 23], which probably reflects the sequence variation among different FG Nups.

**Figure 2.**
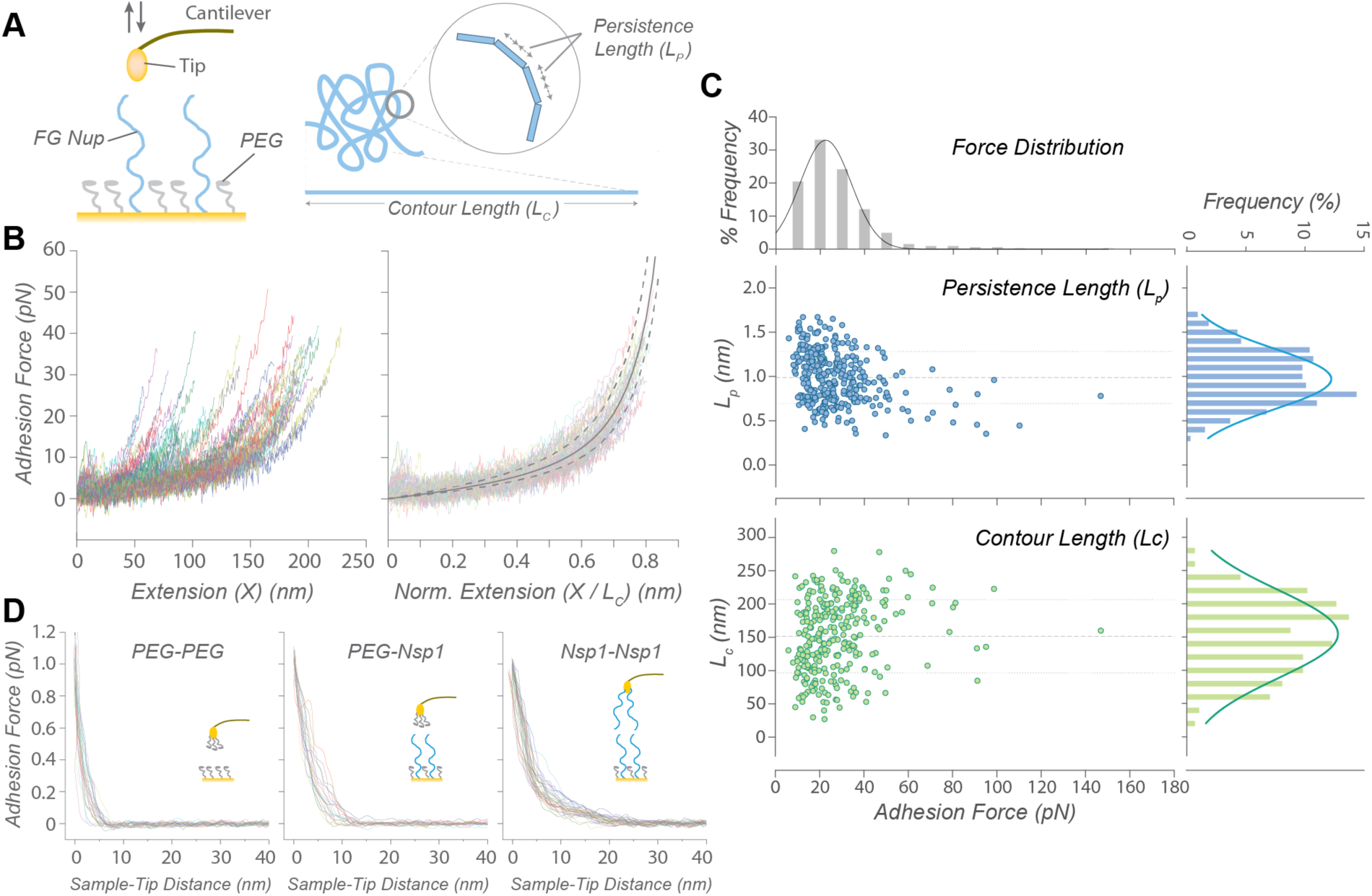
Characterization of an Nsp1FG surface by AFM. (A) Schematic diagram of Nsp1FG approach-pulling experiment and worm-like chain (WLC) model parameters. (B) Representative retraction curves during single Nsp1FG stretching (left), and superimposition of the normalized curves (right). Each curve represents a single stretching event. (C) WLC model estimates of persistence length (*Lp*) and contour length (*Lc*) scatter-plotted against the adhesion force. Their frequency distributions are plotted in respective histograms. (D) Approach curves for layer thickness measurements: Control (PEG-PEG, left), single Nsp1FG layer (PEG-Nsp1, middle), and double Nsp1FG layer (Nsp1-Nsp1, right).

AFM experiments were also performed on a dense brush layer of Nsp1FG, dNsp1FG, to simulate the high density of FG Nups in the NPC [11, 70–72]. The approach (as opposed to retraction experiment above) curves of the FV mapping scans were used to estimate the thickness of the dNsp1FG layer (Fig 2D). When the cantilever tip contacts a stiff surface, a sharp deflection (i.e. ∼90° angle) of the approach curve is expected at the zero sample-tip distance. Since dNsp1FG formed a soft pliable layer on top of the stiff substrate, the cantilever tip sensed a repulsive force while indenting the dNsp1FG layer. This resulted in a curved deflection profile, before the sharp deflection when contacting the stiff sublayer. The onset of the repulsive force was used to estimate the dNsp1FG thickness of ∼10 nm (Fig 2D, PEG-Nsp1). The thickness doubled when dNsp1FG was also on the cantilever (Fig 2D, Nsp1-Nsp1) and was much higher than the hPEG_6_-C_11_-SH control of ∼5 nm (Fig 2D, PEG-PEG). The AFM measurement may have underestimated the layer thickness if: (i) the AFM probe traveled through the top part of the FG Nup layer without a measurable deflection; (ii) the AFM probe sensed a positive deflection only after it compressed the layer; and (iii) the AFM did not reach a hard-contact with the substrate. Nevertheless, the Nsp1-Nsp1 measurement was roughly twice that of PEG-Nsp1 (Fig 2D) and AFM and QCM-D analyses provided similar estimates of the FG layer thickness, suggesting internal consistency of our measurements of the FG-conjugated surface.

### Probing the FG-Kap95 interaction by QCM-D

To enable reproducible analysis from a FG surface, a regeneration procedure that yields maximum removal of remaining analyte at the end of each binding cycle was established. Two macromolecules were used for this optimization. The first, GST-Kap95, dimerizes via GST, providing increased valency for interaction, making it more likely to remain on the FG surface. The second, BSA, has been extensively employed as a test protein to check for surface inertness [73, 74]. Removal of both GST-Kap95 and BSA was achieved using 50 mM HEPES-KOH pH 7.4, 8 M urea, with 0.5% Tween (HUT buffer). While urea can denature the analyte, it is compatible with natively disordered FG constructs, and thus, is an appropriate addition to the regeneration buffer. Indeed, HUT completely regenerated the FG Nup surface (S3 Fig) and thus, was used at the end of every binding experiment.

Once conditions for reversible binding were established, the Nsp1FG surface was tested with different analytes to assess its specificity. Kap95 without the GST domain was able to reversibly bind the surface. Consistent with studies by others [13–16, 25–27], the Nsp1FG surface was essentially inert to non-specific proteins such as BSA and GST, and highly selective for Kap95 (Fig 3A). Thus, the expected qualitative characteristics of the FG-Kap95 interaction were recapitulated by QCM-D. However, further binding analyses revealed that a more detailed interpretation of the data is required.

**Figure 3.**
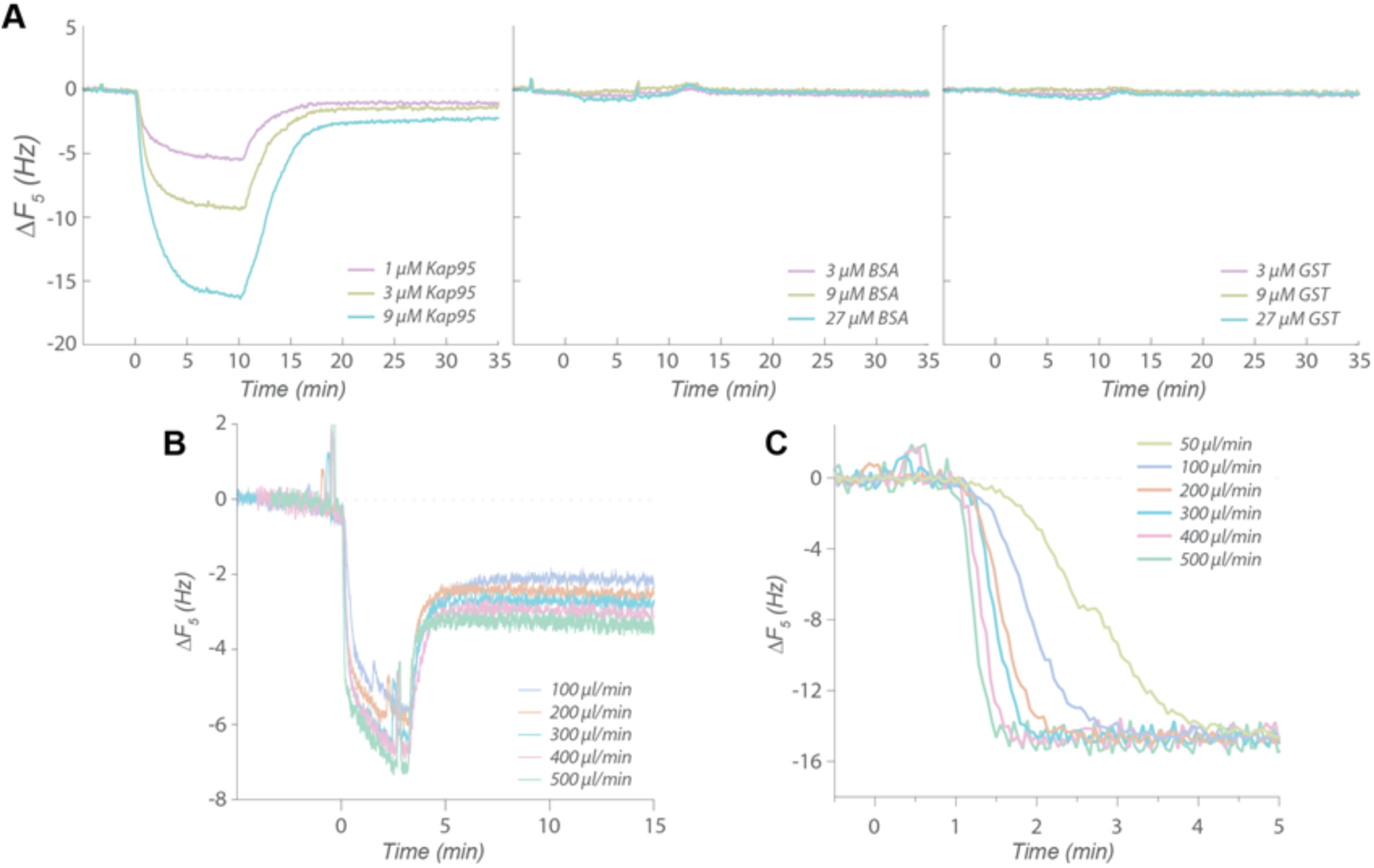
Analyte binding assayed by QCM-D. (A) Binding of different analytes on Nsp1FG surface: Kap95 (left), BSA (middle), and GST (right). (B) Dependence of Kap95 binding to Nsp1FG surface on the flow rate. Changes in resonance frequency (*ΔF_5_*, 5th overtone) upon Kap95 binding are shown for different flow rates. (C) Sensing volume replacement by glycerol solution at different flow rates (see main text). Changes in resonance frequency (*ΔF_5_*, 5th overtone) after introducing feed solution containing glycerol (without any analyte) are shown for different flow rates.

### Influence of mass transport limitation

One of our major concerns was the effect of mass transport limitation, a well-known physical phenomenon among chemical engineers but less frequently examined in the biological literature [35, 36, 75]. Mass transport limitation is observed when the bulk flow over the sensor is insufficiently high relative to the binding kinetics of the protein-protein interaction under investigation to exclude influences of the transport process on the quantitation of kinetic parameters. Bulk flow transports analytes in the feed solution (i.e. mass transport) and the analytes diffuse from the bulk into the volume immediately adjacent to the surface, where they are close enough to interact with the ligands. When the rate of mass transport by bulk flow is sufficiently high, the buildup of analytes on the surface is fast, due to efficient diffusion from the bulk. The rate of protein absorption/binding actually depends on the concentration of the analyte in this region immediately adjacent to the surface, where the analytes are close enough to the surface layer for the interaction to happen, rather than the concentration in the bulk feed solution [35, 36, 75]; thus, when the bulk flow is too low, the concentration of the analyte in the region immediately adjacent to the surface is lower than that in the feed solution because the surface ligand ‘depletes’ the analyte locally, faster than the delivery by the bulk flow. Depletion by the ligand increases with higher ligand density as well as a higher on-rate of the protein-protein interaction. In the dissociation phase of a binding experiment, feed solution is usually a blank solution without any analyte. Under conditions where flow rates are slow and/or the ligand density is high, the escape rate of analytes from the surface can be slower than the ‘rebinding’ rate to the ligand, prolonging the apparent residence time of the analyte on the surface. Therefore, if mass transport limitation is significant but not taken into account, model fitting will yield erroneous kinetic parameter estimates.

There have been numerous studies utilizing surface-based systems to study protein-protein interactions involved in nuclear transport [13–16, 25–27], some of which indeed inferred the existence of mass transport limitations [13, 26, 27]. We initially became concerned about mass transport limitation when we observed that the fraction of GST-Kap95 that dissociates from the surface in our QCM-D system was dependent on the length of the association phase (S10 Fig), i.e. on the saturation level of the surface at the onset of the dissociation phase [35]. Hence, we studied in more detail the effect of flow rate on Kap95 binding alone, which indeed also proved flow-rate-dependent: larger amounts of binding were observed at higher flow rates, indicating mass transport per unit time was limiting at lower flow rates (Fig 3B), and suggesting that there was a lag in the mixing of analyte in the sensing volume.

We suspected that the large sensing volume (∼40 μl) of the QCM-D was a major factor for mass transport limitation. Based on the physical dimension of the reaction chamber, the actual flow rate close to the surface (at 10 nm) is estimated to be 1/10^5^th of the bulk flow, which even at high flow rates remains in the laminar region according to the manufacturer specifications (e.g. average flow velocity of 1.03 mm/s with Reynold’s number of 1.2 at the maximum flow of 500 μl/min). To accurately interpret binding/unbinding kinetics, the sensing volume must be replaced with the feed solution essentially instantaneously. Instead, a mixing lag exists before the concentration of the analyte in the chamber reaches that of the feed solution. Therefore, we measured the extent of mixing lag at various flow rates by replacing an aqueous buffer with one containing glycerol. The higher density of glycerol-containing buffer results in a change in *ΔF*, allowing us to monitor the liquid replacement in real time in the absence of analytes (Fig 3C). A concentration of glycerol of 3% was sufficient to induce a significant change in *ΔF*, yet minimizes the effect of higher density and viscosity. It took ∼30 s to replace the entire sensing volume even at 500 µl/min, implying that only ∼3% of the volume is replaced every second. Such a time scale is many orders of magnitude greater than that of most biochemical reactions, including the FG-TF interactions [17–19, 47].

### Mass transport limitation in SPR experiments

Analogous experiments were also conducted with SPR to test if its smaller sensing volume would allow measurement of the same FG-TF interaction without mass transport limitation. Although the same conjugation protocol used for QCM-D could not be implemented due to the conjugation chemistry available for the SPR system (XPR36, BioRad), the conditions for the binding experiments were optimized to minimize non-specific binding to a bare surface passivated with beta-mercaptoethanol (S11 Fig). A segment of Nsp1FG that contains six repeats of the FxFG-type region (FSFG_6_) was used for SPR. The binding of analytes to FSFG_6_ was consistent with our QCM-D experiments with Nsp1FG and FSFG_6_ (see below), and also with previously reported SPR results [14–16]. We observed Kap95 to bind FSFG_6_ reversibly, whereas non-specific proteins did not exhibit any significant binding (Fig 4A), suggesting that the SPR measurements were consistent with the QCM-D setup.

**Figure 4.**
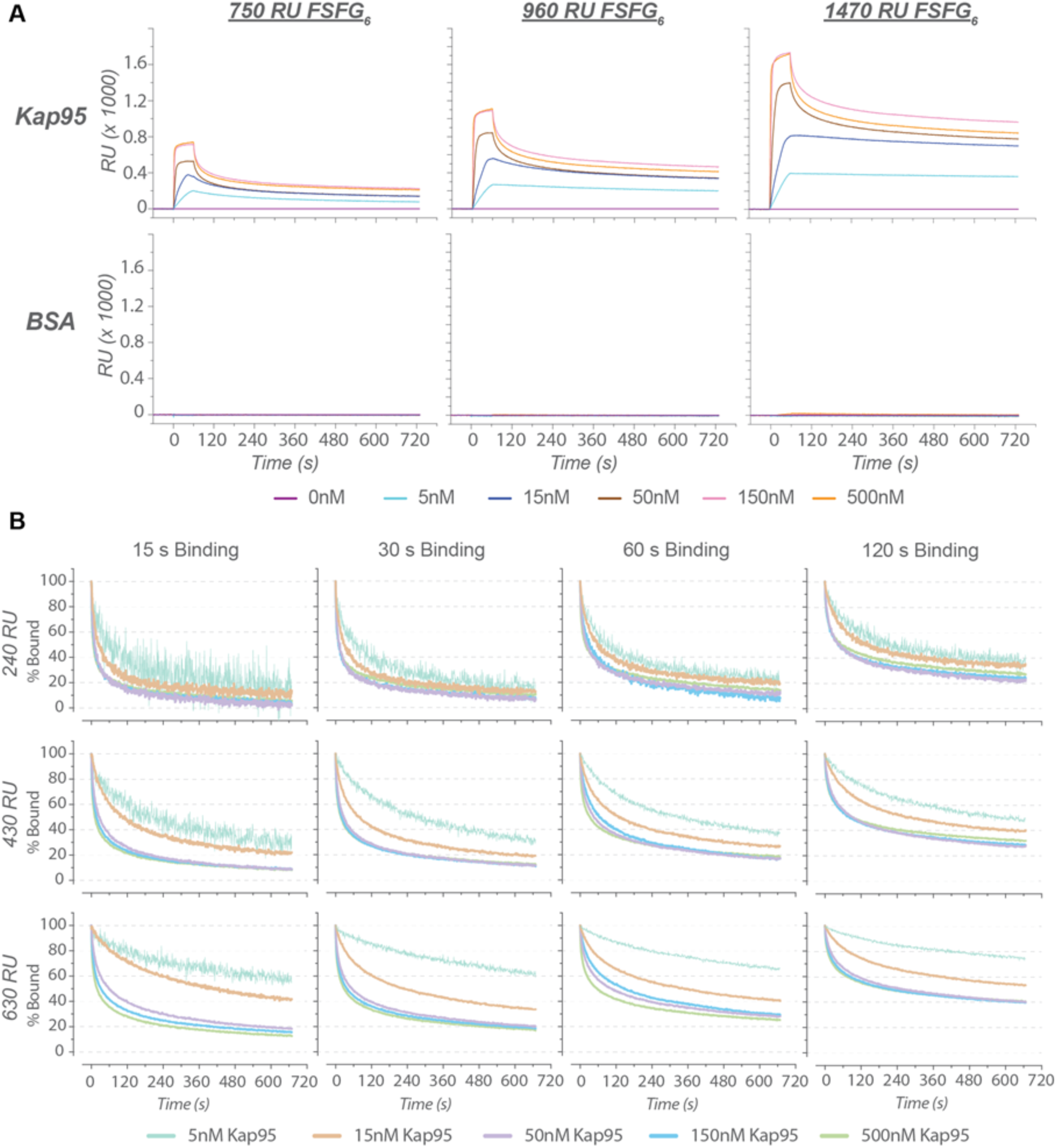
Analyte binding assayed by SPR. (A) Kap95 (top row) and BSA (bottom row) binding to a FSFG6 surface at different surface ligand densities. RU (response unit) values indicate the amount of FG protein on the surface: low (750 RU), medium (960 RU), and high (1470 RU), respectively. (B) Normalized dissociation curves of Kap95 to assess the existence of mass transport limitation. Different lengths of the association phase (columns) and different ligand densities (rows) are shown. Colors indicate different Kap95 concentrations. Respective unnormalized curves including the association phase are shown in S12 Fig.

To test for the existence of mass transport limitation, we first analyzed the SPR sensorgrams. In the absence of mass transport limitation, all the normalized dissociation curves from different analyte concentrations should converge to a single exponential decay. However, normalized Kap95 dissociation curves deviated from a single exponential decay, indicating the existence of more than simple binding events (Fig 4B). The characteristic signatures of mass transport limitation [35] were present, as evidenced by: (1) the curvature of the normalized dissociation curves is dependent on Kap95 concentration, and steeper at higher Kap95 concentrations; (2) the relative amount of Kap95 retained on the surface after a fixed time period increases with the density of the surface ligand; and (3) the relative amount of Kap95 retained on a surface with constant ligand density increases with the amount of Kap95 bound to the surface (i.e. extent of surface saturation) at the onset of the dissociation phase. These observations suggest that mass transport limitation is also present in the SPR experiments. Our numerical simulations also suggest that mass transport limitations can be induced for even simpler interactions in this type of SPR chamber (S2 Text, S13 Fig, and S2 Table).

We next assessed whether the SPR data can be interpreted by commonly used kinetic models and by a simple Langmuir isotherm. Fitting results to those models are summarized in S14 Fig and S2 Table. A simple pseudo-first order kinetic model did not fit well to the experimental binding curves, as indicated by the large 𝜒^2^. Inclusion of the mass transport limitation term also did not improve fitting to a one-to-one model. Fitting was better for more complex models, specifically for the two-state model, where conformational change of the ligand-analyte complex is assumed, and for the heterogeneous ligand model, where two types of independent ligands are assumed. In contrast, the heterogeneous analyte model, where two types of independent analytes were assumed, did not fit well. The goodness of fit for each model tested here can be assessed visually in S14 Fig (see also S2 Table). The data could be fitted equally well by the two-state model and the heterogeneous ligand model, suggesting that the interpretation of the data depends on an arbitrary choice of the model in the absence of independently obtained mechanical information. The trend in our fitting results is similar to that found in previous works, where more complex models, including a Hill model [13, 25–27], yielded better fits to QCM-D binding curves than a simple Langmuir isotherm.

### Existence of more than one binding mode

In addition to the curve fitting analysis, further experiments were conducted to assess the biochemical relevance of our binding data by utilizing a mutant variant of FG constructs where the phenylalanines in FSFG constructs were replaced with serines (i.e. SSSG motifs). Structural study indicate that FG Nups interact with karyopherins such as Kap95 by inserting the hydrophobic sidechain of their phenylalanine residues into the hydrophobic pockets of karyopherins, underscoring the requirement of the phenyl group in this interaction [43]. This Phe > Ser mutation was demonstrated to abolish interactions with TFs [19, 46]. Thus, a six-repeats construct, SSSG_6_, was tested to determine if this total loss of interaction is observed in QCM-D and SPR experiments.

According to the QCM-D measurements, FSFG_6_ and SSSG_6_ layers behaved similarly, as they formed rigid layers due to their short length (S15 Fig), unlike the viscoelastic Nsp1FG layer (Figs 1C and S4). As expected, Kap95 bound strongly to FSFG_6_, though to a lesser extent than to Nsp1FG due to the reduced number of FG motifs (Fig 5A). More interestingly, Kap95 also interacted significantly with the surface-bound SSSG_6_ construct, albeit to a lesser extent than the Phe counterpart, but greater in comparison to the binding pattern of non-specific analytes to Nsp1FG (Fig 4A). Similar observations were made in the SPR experiments: Kap95 binds to the SSSG_6_ construct to a significant extent (Fig 5B), suggesting that Kap95 can be retained on SSSG_6_ surface despite the lack of FG motifs to mediate any specific binding. However, NMR data show that linker residues of FG repeat regions do not participate in the interactions with TFs [17, 18], and Phe > Ser mutations in Nsp1FG abolish interaction with TFs *in vivo* [76], such that the SSSG_6_ construct should not bind TFs. Since Kap95 did not bind to a bare SPR sensor under our experimental conditions (S11 Fig), we conclude that the Kap95 interactions are a result of unexpected binding to the SSSG_6_-conjugated surface, further complicating the measurements in conjunction with the influence of mass transport limitation.

**Figure 5.**
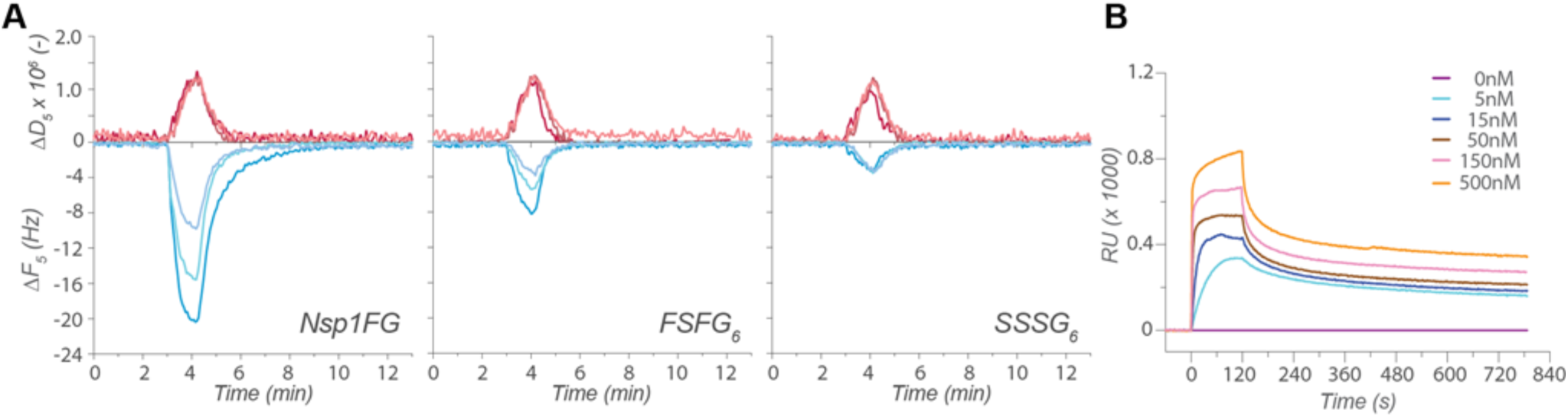
Analyte binding experiments with SSSG_6_ mutant variant. (A) Kap95 binding on Nsp1FG (left), FSFG6 (middle), and SSSG_6_ (right) monitored by QCM-D. Changes in resonance frequency (*ΔF_5_*) (bottom, blue) and energy dissipation (*ΔD_5_*) (top, red) for the 5^th^ overtone are shown. Kap95 concentrations are indicated by different shades of each color: light, medium, and dark shades indicate 1, 3, and 9 µM Kap95, respectively. (B) Kap95 binding on SSSG_6_ surface (∼920 response units (RUs) of SSSG_6_ conjugated) by SPR. Kap95 concentrations are indicated by different colors.

## Discussion

FG Nup constructs and their interactions with Kap95 were systematically characterized by AFM, QCM-D, and SPR. Nsp1FG grafting conditions were optimized and cross-validated by AFM and QCM-D, which yielded a consistent picture on the viscoelastic behavior of Nsp1FG. Both QCM-D and SPR demonstrated the *apparent* specificity of the FG-grafted surfaces to the TF, Kap95. Therefore, highly robust experimental systems were set up to reproducibly collect data on TF binding to FG-grafted surfaces. The results from AFM, QCM-D, and SPR were in agreement with each other and with previous observations in the literature. However, our further investigations demonstrated the existence of mass transport limitation and extra binding mode(s). The existence of mass transport limitations has been acknowledged in the nuclear transport field [13, 26, 27]. The present work demonstrates the importance of taking mass transport limitations properly into account, as well as of considering unexplained physical phenomena that are implicit in the binding data. One could attempt to infer microscopic mechanisms that produced the binding curves, however, such deconvolution processes require assumptions about those mechanisms (e.g. what processes exist and how they depend on each other, along with all their parameters). While there have been attempts to model the generative process of the surface binding data, we and others found that the underlying dynamics of the “fuzzy” FG-TF interactions to be quite complex [17–19]. Since it is quite difficult to validate the assumptions we would have to make to model the surface binding data, we opted out from making such inferences in this work.

### Complexities of FG-TF binding reactions on a surface

Mass transport limitation is a significant but often overlooked issue in the field of SPR and QCM-D [35]. Although we observed that our surface-bound FG layers were selective for Kap95, the binding reaction appeared to be affected by mass transport limitation with both QCM-D and SPR. The likely contributing causes to this mass transport limitation include: (i) the physical scale of the flow systems, introducing a mixing lag, (ii) fast kinetics of FG-TF interactions, and (iii) the high multivalency of TFs and FG Nups that facilitates rebinding. The charge properties of the interactors may also affect their bulk mass transport behavior. The dimension of the QCM-D sensor is relatively large (millimeter ∼ centimeter range), so that it takes ∼30 s to replace the entire sensing volume of the chamber (Fig 3C), which resulted in the flow-rate-dependent binding of Kap95 (Fig 3B). In an attempt to alleviate the effect of mass transport limitation, we used SPR to study FG-TF interaction; its sensing volume is four orders of magnitude smaller than that of QCM-D chamber. However, our SPR experiments still appeared to suffer from a significant mass transport limitation (Fig 4). In addition, the significant interaction of SSSG_6_ with Kap95 suggests the existence of an FG-independent binding mode, further complicating the analysis of the data (Fig 5).

Although our results do not necessarily invalidate SPR and QCM-D for studying the physical properties of such surface layers, they indicate that data garnered using these methods must be interpreted with extreme care when studying a complex interaction, like the one here with rapidly interacting proteins, when the surface density of ligand binding sites are high, thick and viscous. This is especially relevant in studying IDPs. Indeed, FG-TF interaction dynamics at the molecular level were shown to be even faster than the ∼milliseconds time scale of NMR experiments [19], with association rate constant estimated to be ∼1.5·10^9^ M^−1^·s^−1^ [18]. Thus, the length scale and time-scale of the SPR and QCM-D are too large to resolve any of the underlying molecular events considered here. Furthermore, the models tested with data from SPR largely underestimates the fastest component; the fastest rate estimated from the kinetic model fits was ∼3.0·10^6^ M^−1^·s^−1^ (S2 Table), a rate that is orders of magnitude slower than the molecular rate. We considered equilibrium analysis because of its time-independence. However, it proved difficult to establish if binding even reached an equilibrium with QCM-D, as the deposition of analytes continued to bind on a time-scale of minutes (S10 Fig). Moreover, our data suggests the existence of multiple types of binding events, and thus, any estimation of dissociation constant(s) would be heavily model-dependent, underscoring the limitation of extracting mechanistic information of a complex binding reaction solely from SPR and QCM-D sensorgrams. Thus, we did not attempt to draw any quantitative conclusions from these data regarding the mechanism of nuclear transport. In addition, previously reported high affinities for FG-TF interactions from surface-based methods [8–16, 20] require re-evaluation in this light. The reported *K_D_*s in those studies reflect the ‘residence time’ of TFs in FG layers (see below) rather than the thermodynamics of the binding mechanism at the molecular level. We recently emphasized the distinction between the global residence time of a protein in a loose protein-protein interaction (i.e. how long the two molecules stay associated with each other) and the half-life of an individual per-site interaction (i.e. how long an *individual bond* involved in the interaction persists) [17–19].

### Phenomenological *K_D_*s and microscopic mechanisms

Following the discussion above, we argue that it is not possible to accurately obtain microscopic *K_D_* (s) for a complex binding reaction(s) simply based on a sensorgram without knowing the exact mechanism of binding on the surface-bound layer. The use of simple Langmuir isotherms to describe such complex reactions only yields ‘phenomenological (or apparent) *K_D_*s’, which may be useful for estimating the partitioning of analytes between bulk flow and surface layer but do not provide precise thermodynamic information about the binding mechanism. Indeed, in the nuclear transport field, Richter and others have noted that their FG-TF binding data from QCM-D are “consistent with the presence of a spectrum of binding sites that cover a range of affinities,” suggesting the phenomenological nature of *K_D_*s derived from such analytical models [13, 27]. For a complex protein-protein interaction, it is generally very difficult to accurately determine (i) the number of different binding mechanisms, (ii) the dependencies of such mechanisms on each other, (iii) the relative contributions to the overall binding curve from each mechanism, and (iv) their individual kinetic and thermodynamic parameters. A simple Langmuir isotherm, or even the simplest form of Hill equations that is commonly used, would not properly represent such complex interactions [77]. Our experiments indicate that there are at least three kinds of physical events that contribute to the sensorgrams: FG-dependent binding, FG-independent binding, and mass transport limitation. In a case like this, where mass transport limitation is further complicated by multivalency, it is also difficult to estimate the extent of rebinding events within the surface layer because it is not manifest in the sensorgram. For instance, it is not possible to distinguish species with a long half-life of complexation from those that are simply retained on the surface by way of frequent rebinding. If the model does not properly account for the intricacies of multivalency in the presence of mass transport limitation, it would appear as if the binding is of high affinity with slow dissociation rates, leading to a misinterpretation of the data. Thus, phenomenological *K_D_*s can be useful in describing macroscopic behaviors, but inferring their underlying mechanisms and their corresponding kinetic and thermodynamic parameters requires a set of assumptions; hence, soundness of such models fundamentally depends on the validity of the assumptions they make, regardless of their mathematical sophistication [78].

This is particularly relevant for analyzing IDPs, such as FG Nups. IDPs are known to have a broad spectrum of behaviors compared to folded proteins. The conformational space they can explore is immense and the mechanisms of interactions with their binding partners are diverse and complex, many of which are very difficult to track. For example, some IDPs bind to their partners by induced folding, while others maintain their intrinsic disorder in “fuzzy” interactions, like FG Nups [47, 79]. The mechanisms of these interactions are quite complex and the interaction mixture consists of ensembles of different configurational states, as suggested in our recent study [19]. Therefore, lacking the capability to finely resolve the mechanistic aspects of protein-protein interactions (e.g. distinct energy levels, defined interaction surfaces), the use of discrete analytical models on SPR and QCM-D data is probably not suitable for studying the mechanisms of IDP interactions. One exception are cases where QCM-D and AFM are used to study the morphological and viscoelastic features of a bulk IDP layer.

### Macroscopic instruments versus nanoscopic biological systems

The major cause of mass transport limitation in this study appeared to be the length- and time-scale of the experimental setup. Dimensions of QCM-D and SPR sensing volumes are orders of magnitude larger than the biological machineries they are exploring, with typical scales of tens of nanometers. For example, the FG-Nup filled central tube of the NPC spans ∼50 nm, and thus is at least four orders of magnitude smaller than the above instruments. Under such nanoscopic environments, mass transport limitation would still exist, but its effect would scale similarly to those of molecular interactions and diffusion. However, under macroscopic systems such as SPR and QCM-D, mass transport limitation magnifies with the physical dimensions of the system (e.g. total number of binding sites increases with scale). The number of analyte binding/unbinding cycles it takes to randomly escape from the FG milieu is small at the nanometer scale. In contrast, the FG surface layers on SPR and QCM span microns to millimeters, so the number of on-off cycles to escape can be quite large (with a commensurate substantially longer residence time). Thus, the macroscopic dimensions of the instruments could well overshadow the molecular interactions and diffusion effects that are biologically relevant at nanoscopic scales. Solution studies *in vitro*, such as ITC, stopped-flow kinetics, NMR, and appropriately-scaled NPC mimics such as nanoscopic pores [48, 80–82] and DNA origami pores [83, 84] offer the possibility of studying FG-TF interactions at a scale more commensurate with that of nuclear transport *in vivo*.

## Acknowledgements

We thank the High Throughput and Spectroscopy Resource Center at the Rockefeller University. We thank Drs. Roderick YH Lim, Larisa E. Kapinos, Ralph P Richter, Bart W Hoogenboom, David Cowburn, and Samuel Sparks for their helpful comments on the manuscript.

This work was supported by National Institutes of Health grants P41GM109824, R01GM112108 and R01GM117212 (to M.P. Rout)

