## supplementary_information for "Interactions of nuclear transport factors and surface-conjugated FG nucleoporins: Insights and limitations"

**Supplementary Text 1. Primary sequences of the protein constructs**

**Nsp1FG-Cys-$His_6$**
MSTGAGAFGTGQSTFGFNNSAPNNTNNANSSITPAFGSNNTGNTAFGNSNPTSNVFGSNNST
TNTFGSNSAGTSLFGSSSAQQTKSNGTAGGNTFGSSSLFNNSTNSNTTKPAFGGLNFGGGNN
TTPSSTGNANTSNNLFGATANANKPAFSFGATTNDDKKTEPDKPAFSFNSSVGNKTDAQAPTT
GFSFGSQLGGNKTVNEAAKPSLSFGSGSAGANPAGASQPEPTTNEPAKPALSFGTATSDNKT
TNTTPSFSFGAKSDENKAGATSKPAFSFGAKPEEKKDDNSSKPAFSFGAKSNEDKQDGTAKP
AFSFGAKPAEKNNNETSKPAFSFGAKSDEKKDGDASKPAFSFGAKPDENKASATSKPAFSFGA
KPEEKKDDNSSKPAFSFGAKSNEDKQDGTAKPAFSFGAKPAEKNNNETSKPAFSFGAKSDEK
KDGDASKPAFSFGAKSDEKKDSDSSKPAFSFGTKSNEKKDSGSSKPAFSFGAKPDEKKNDEV
SKPAFSFGAKANEKKESDESKSAFSFGSKPTGKEEGDGAKAAISFGAKPEEQKSSDTSKPAFT
FGCLEHHHHHH*

**$FSFG_6$-Cys-$His_6$**
MNETSKPAFSFGAKSDEKKDGDASKPAFSFGAKPDENKASATSKPAFSFGAKPEEKKDDNSS
KPAFSFGAKSNEDKQDGTAKPAFSFGAKPAEKNNNETSKPAFSFGAKSDEKKDGDASKPACL
EHHHHHH*

**$SSSG_6$-Cys-$His_6$**
MNETSKPASSSGAKSDEKKDGDASKPASSSGAKPDENKASATSKPASSSGAKPEEKKDDNSS
KPASSSGAKSNEDKQDGTAKPASSSGAKPAEKNNNETSKPASSSGAKSDEKKDGDASKPACL
EHHHHHH*

**Supplementary Text 2. Simulating simple protein-protein interactions by multiphysics finite element model.**

To test the prevalence of mass transport limitation, we conducted numerical simulations on hypothetical protein-protein interactions in a simulated SPR chamber. For these simulations, we modeled a set of proteins with lower molecular weights than those of the FG constructs and Kap95 used in our experiments. These lower molecular weight proteins diffuse faster and should thus provide useful lower limits on the magnitude of mass transport limitations in our experiments. To numerically simulate such interactions, a multiphysics finite element model was developed to understand the role of convection mass transfer in a particular geometry of the SPR chamber used in the experiment. The SPR chamber was set to be a rectangular prism 1 mm deep, 1 mm wide, and 0.5 mm high. The surface ligand density, molecular weight of the ligand, molecular weight and concentration of the analyte, and finally binding site kinetics used in the model are summarized in Table S2. The model considered several aspects of the system. The volumetric flow rate of 10, 100, and 1,000 μl/min were used as the inlet boundary condition for the flow, which all resulted in laminar flow field. Transport through the SPR system was modeled with the following system of equations and boundary conditions, assuming the analyte solution was dilute:

$$\frac{dc_i}{dt} + \nabla \cdot (-D_i \nabla c_i) + \boldsymbol{u} \cdot \nabla c_i = R_i$$

$$\boldsymbol{N}_i = -D_i \nabla c_i + \boldsymbol{u} c_i$$

Boundary condition at the wall: $-\boldsymbol{n} \cdot \boldsymbol{N}_i = 0$

Boundary condition at the inlet: $\boldsymbol{n} \cdot \boldsymbol{N}_i = \boldsymbol{n} \cdot (\boldsymbol{u} c_{0,i})$

Boundary condition at the outlet: $-\boldsymbol{n} \cdot D_i \nabla c_i = 0$

where $c_i$ is the concentration of species *i*, $c_{0,i}$ is the concentration of species *i* at the inlet (i.e., feed solution concentration), $D_i$ the diffusivity of species *i*, $\boldsymbol{u}$ the velocity, $R_i$ the source term representing the net rate of formation of species *i* per unit volume by chemical reaction (i.e. binding/unbinding using Langmuir model), $N_i$ the molar flux of species *i*. The diffusivity of the analyte molecules themselves, $D_{analyte}$, were calculated from their molecular weights using Polson equation:

$$D_{analyte} = \frac{9.40 * 10^{-15} * T}{\mu * \sqrt[3]{MW_{protein}}}$$

where *T* is the temperature, *μ* the solution viscosity, and *MW* the molecular weight of the analyte. As shown by the equations above, transport of analytes in the bulk solution was coupled with a surface reaction, $R_i$, on the lower surface of the SPR chamber that represented analyte binding to its ligand, using a Langmuir binding model.

In a simulation, the SPR chamber was devoid of analytes initially and the solution containing the analyte was introduced into the chamber at $t$ = 0 s. Transient flow simulations were run until the outlet concentration of the analyte reached a steady-state value equal to the inlet concentration. The outlet analyte concentration (averaged over the exit 1 mm x 0.5 mm surface area) and bound surface analyte amount (bound on the lower 1 mm x 1 mm SPR reaction area) were charted to see how convection, diffusion, and the binding reaction all interact to govern the surface binding in the SPR instrument.

**Figure S1. QCM-D - Comparison of gold and silica sensor.**

Nsp1FG was conjugated via its C-terminal cysteine to silica or gold surface without (A, C) or with mPEG passivation (B, D), respectively. The data shown are from 15 min binding-unbinding experiments to test the inertness of the two different surfaces. 1 µM Kap95 was used in all experiments.

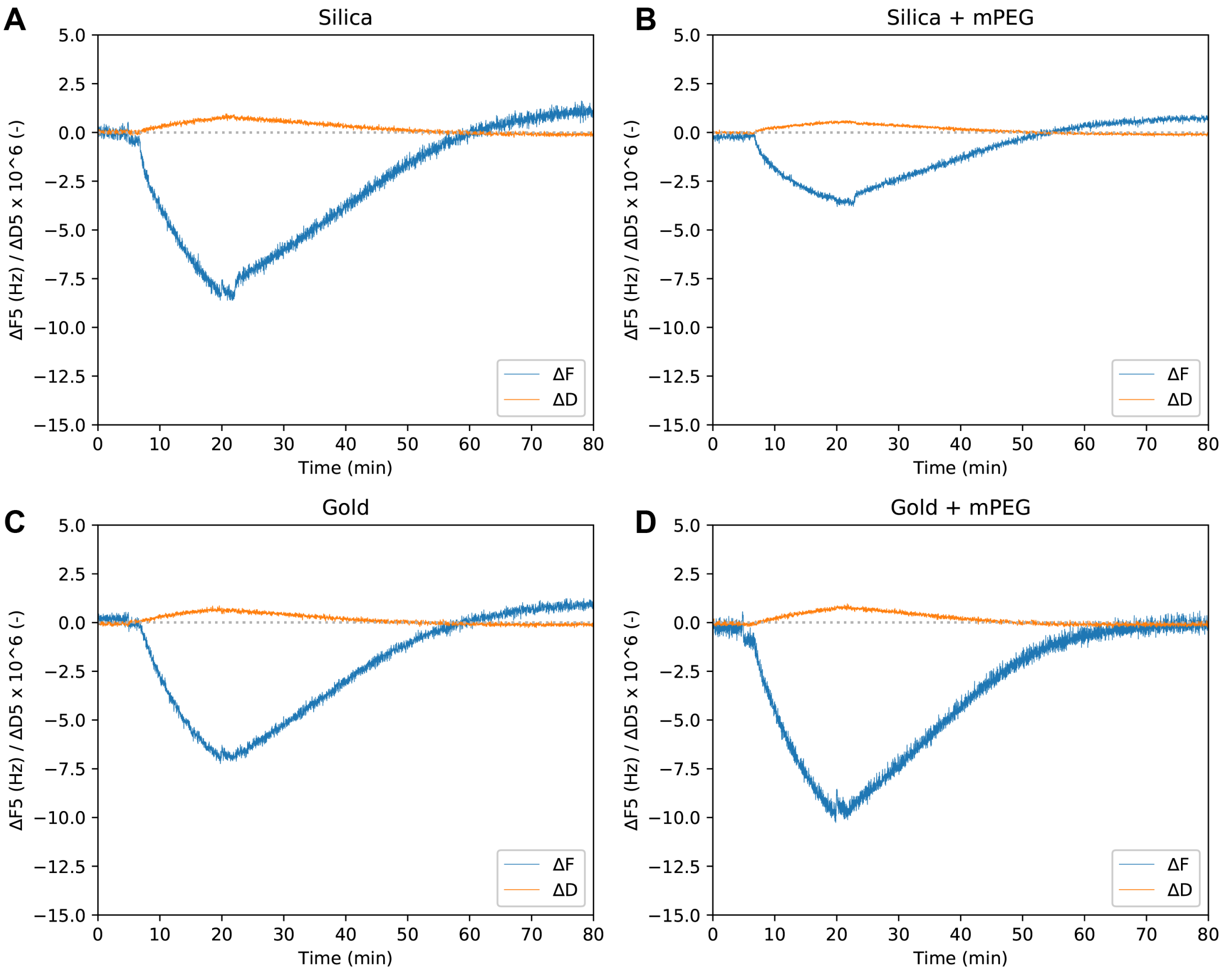

**Figure S2. QCM-D - Comparison of PEG passivators.**

Gold sensors were modified with 2 mM solutions of (A) HS-$(CH_2)_{11}$-$EG_3$-$OCH_3$ ($mPEG_3$), (B) HS-$(CH_2)_{11}$-$EG_3$-OH ($hPEG_3$), and (C) HS-$(CH_2)_{11}$-$EG_6$-OH ($hPEG_6$) in ethanol, overnight and at room temperature. The data show a 30 min binding-unbinding experiment using a 300 µM BSA solution, in order to test the inertness of the different passivated surfaces. $\Delta F$ and $\Delta D$ indicate the change in resonance frequency and dissipation for the 5th overtone, respectively.

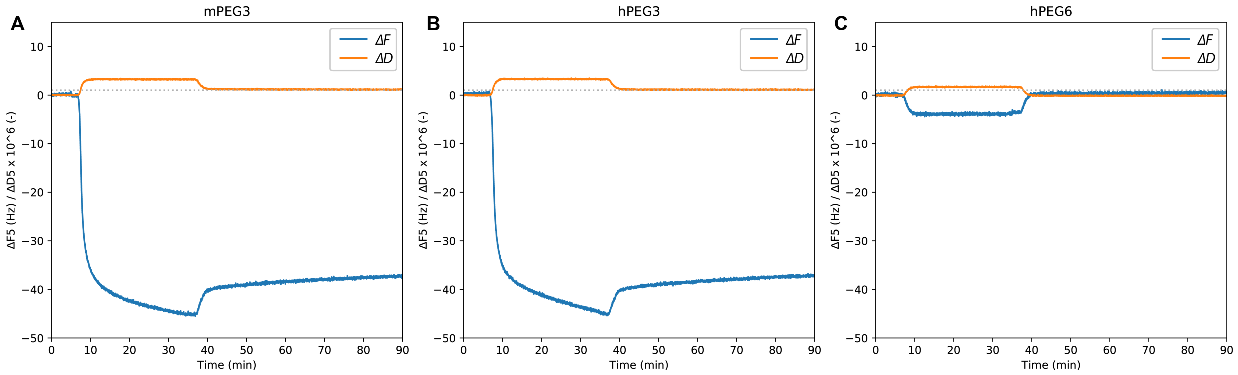

**Figure S3. QCM-D - Surface regeneration.**

(A) Two silica sensors were modified with mPEG (upper) and Nsp1FG (lower) and then tested with GST-Kap95. The data show 6 experiments performed serially, varying binding time and buffer condition: (i) 2 min in TB-D, (ii) 2 min in TBS-D, (iii) 2 min in TBT-D, (iv) 60 min in TB-D, (v) 60 min in TBS-D, (vi) 60 min in TBT-D. A HUT wash step was included between each experiment to regenerate the surface. (B) Magnified views of the HUT washing steps in (A). (C) Multiple 1μM Kap95 binding experiments were repeated on the same Nsp1FG sensor. Regeneration with HUT solution and reproducibility of Kap95 binding was tested. Each color indicates replicate binding events with surface regenerations in between. F5 and D5 indicate 5$^{th}$ overtone of $\Delta F$ and $\Delta D$, respectively. TB: 20 mM HEPES-KOH (pH 7.4), 110 mM KOAc, 2 mM $MgCl_2$, 10 μM $CaCl_2$, and 10 μM $ZnCl_2$; TB-D: TB with 5 mM DTT; TBS-D: TB-D with 150 mM NaCl; TBT: TB with 0.1% v/v Tween20; TBT-D: TBT with 5 mM DTT; TBT-PVP: TBT with 0.3% (w/v) polyvinylpyrrolidone. HUT: 50 mM HEPES, 8 M urea and 0.5% Tween20.

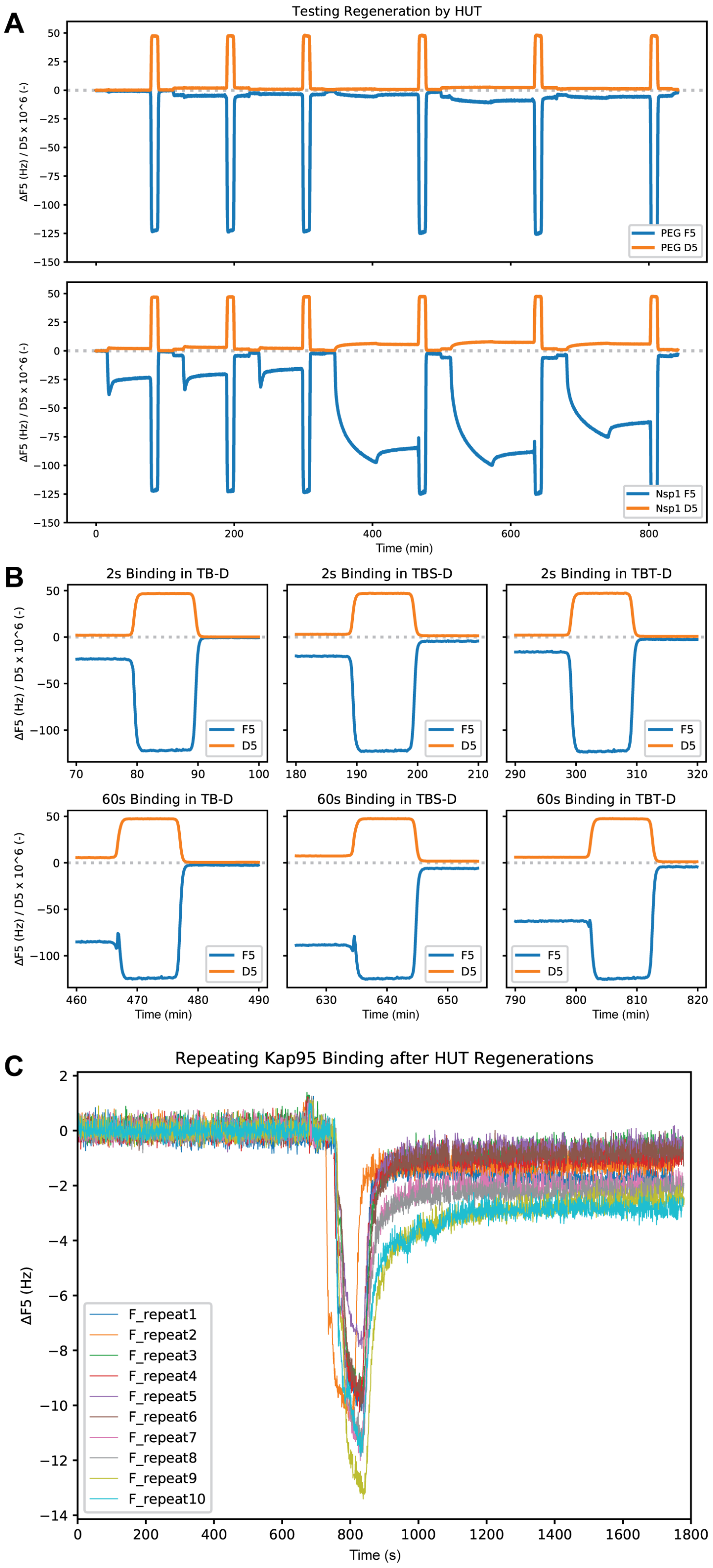
A
Testing Regeneration by HUT
ΔF5 (Hz) / D5 x 10^6 (-)
PEG F5
PEG D5
Nsp1 F5
Nsp1 D5
Time (min)
B
2s Binding in TB-D
2s Binding in TBS-D
2s Binding in TBT-D
60s Binding in TB-D
60s Binding in TBS-D
60s Binding in TBT-D
F5
D5
Time (min)
C
Repeating Kap95 Binding after HUT Regenerations
ΔF5 (Hz)
F_repeat1
F_repeat2
F_repeat3
F_repeat4
F_repeat5
F_repeat6
F_repeat7
F_repeat8
F_repeat9
F_repeat10
Time (s)

**Figure S4. QCM-D - Different harmonics during Nsp1FG conjugation.**

The changes in frequency ($\Delta F$) and energy dissipation ($\Delta D$) for several overtones ($3^{rd}$, $5^{th}$, $7^{th}$, $9^{th}$, and $11^{th}$) recorded in real time during Nsp1FG conjugation are shown. The Nsp1FG conjugation (60 min + 10 min buffer wash) was followed by a HUT wash (5 min), two passivation steps with mPEG (60 + 10 min buffer wash) and beta-mercaptoethanol (βME) (60 + 10 min buffer wash), and a final HUT wash (5 min). The spreading of the different harmonics indicates the viscoelastic nature of the Nsp1FG layer coupled to the sensor surface.

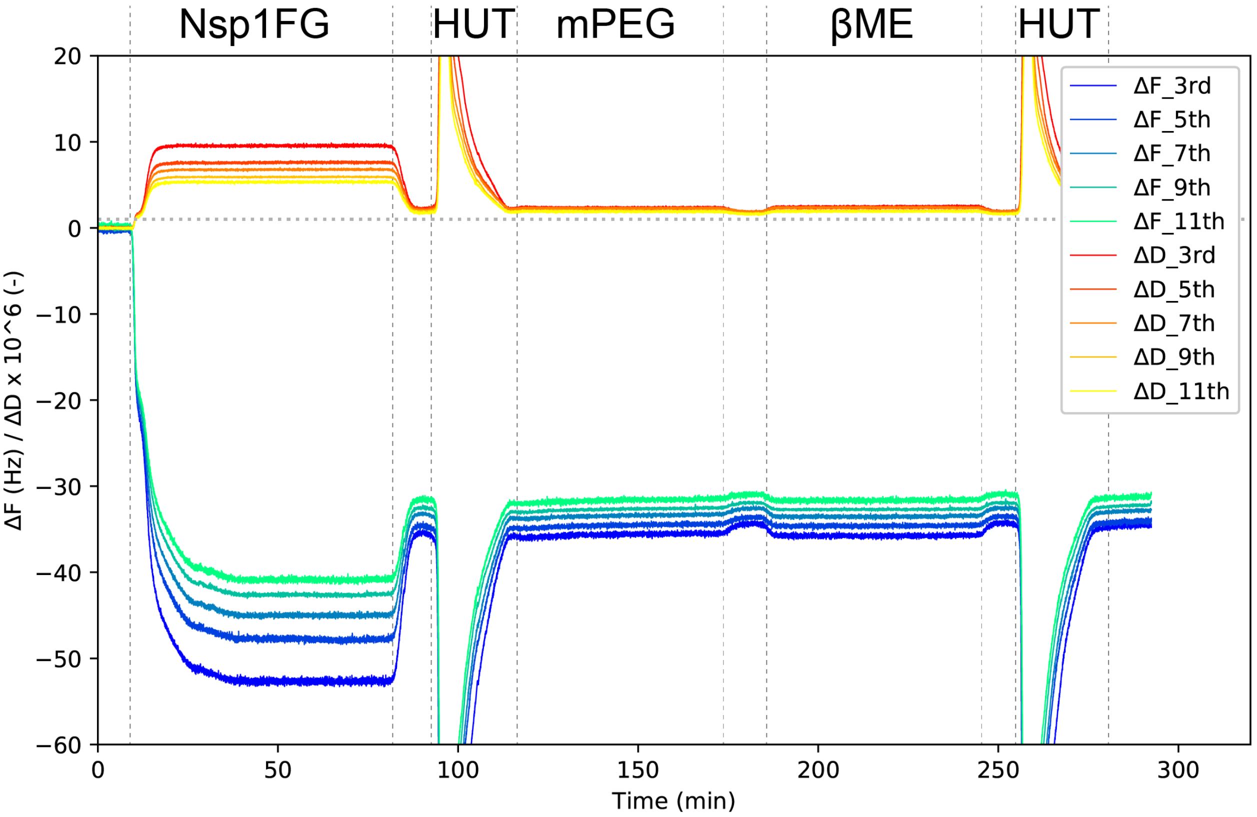

**Figure S5. AFM (volume force mapping) - Surface passivation with $hPEG_6$-$C_{11}$-SH.**

Single molecule force spectroscopy maps between a gold-coated AFM tip and gold surfaces (A) without any passivation and after passivation with hPEG for, (B) 5 or (C) 30 minutes. White pixels (as the ones selected by blue circles in A-C) indicate strong gold/gold interactions (lack of passivation), while dark pixels (as the ones selected by red circles in B-C) show small or no adhesion (~passivated surface). (A) Without passivation, strong non-specific interactions (i.e. few hundreds of pN) between the AFM tip and the substrate resulted during volume force mapping, as indicated by the numerous high adhesions events (i.e. white pixels). The non-homogeneous nature of this interaction indicated by the spread of light and dark pixels, corresponding to stronger and weaker interactions, respectively, was likely due to damage or contamination on the AFM cantilever tip during the scan. (B) A 5 minutes incubation of the gold substrate with $hPEG_6$-$C_{11}$-SH allowed almost complete passivation, resulting in a mostly (~90%) inert surface with weak or no adhesion between the AFM cantilever tip and the gold surface: The average adhesion force, $F_{ad}$ was less than 5 pN. Sparse spots resulted in high adhesion events similar to the negative control, suggesting that bare gold was still available for subsequent Nsp1FG immobilization. (C) A 30 minutes $hPEG_6$-$C_{11}$-SH incubation did not significantly differ from the 5 minute $hPEG_6$-$C_{11}$-SH incubation, which was expected since complete passivation of the surface (i.e. formation of a self-assembled monolayer due to the 11 carbons in the alkane chain, $C_{11}$) requires few hours of incubation (see reference [61] in main text). Force-distance curves for (D) gold-gold adhesion and (E) no adhesion. (F) Histogram representation of $F_{ad}$ for the 5 min sample SMFS-map showed almost complete passivation, with $F_{ad}$ < 5 pN.

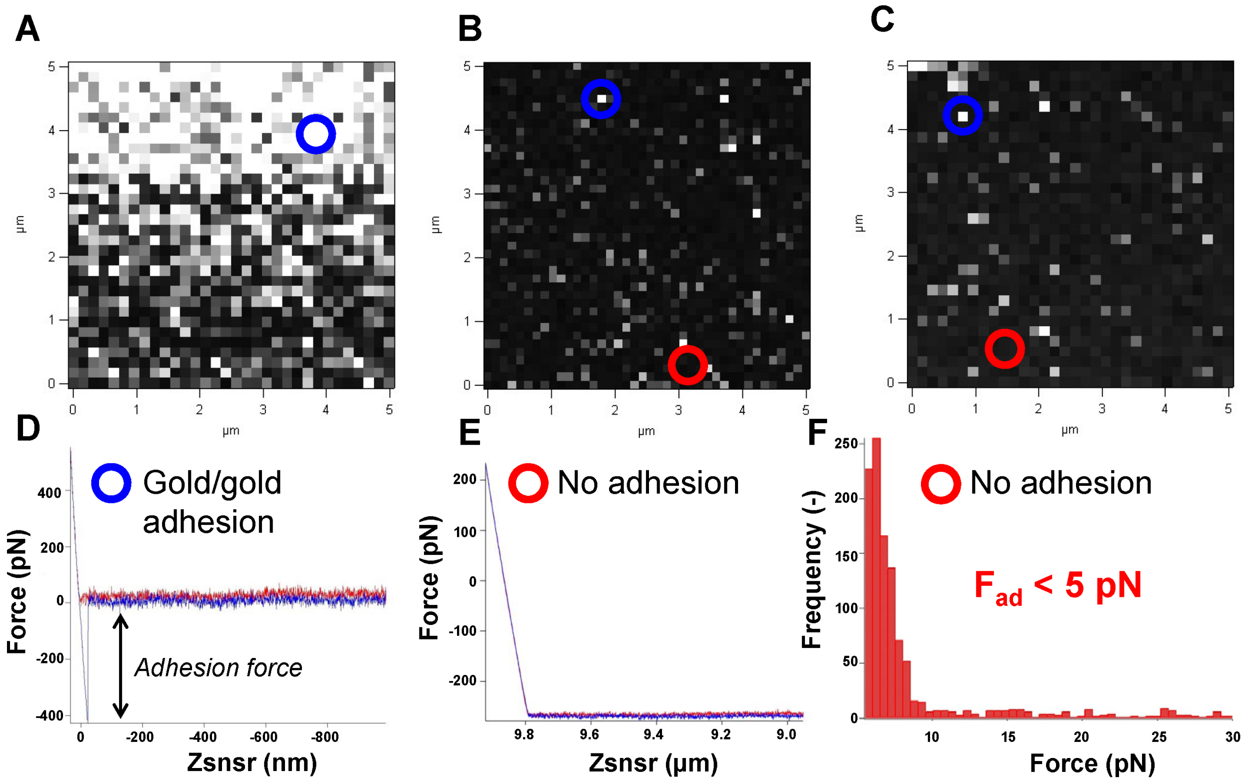

**Figure S6. AFM - Force-map of Nsp1FG for single molecule stretching.**

(A) Single molecule force spectroscopy map of sNsp1 sample. Force-distance curves for (B) no adhesion (dark pixels in (A). An example indicated by a red circle) and (C) adhesion (white pixels in (A). An example indicated by a blue circle). Most of the surface exhibited no binding/adhesion to the AFM cantilever tips, as expected from the passivation. The adhesion events showed different characteristic patterns: (i) single-molecule stretching without an initial adhesion peak (74 out 1,600 ~5%) or (ii) with an initial adhesion peak (64 out 1,600 ~4%), (iii) multiple stretching events (20 out 1,600 ~1%) or (iv) non-reproducible surface/tip interactions (most likely due to non-specific/background interactions – data not used). A data set consisted of an array of 1,600 (40 x 40) force measurements, scanning in contact mode at 1 μm/s an area of 5 x 5 $\mu m^2$, with each pixel point spanning an approximate width of 125 nm in both X and Y. The trigger force was set to 500 pN.

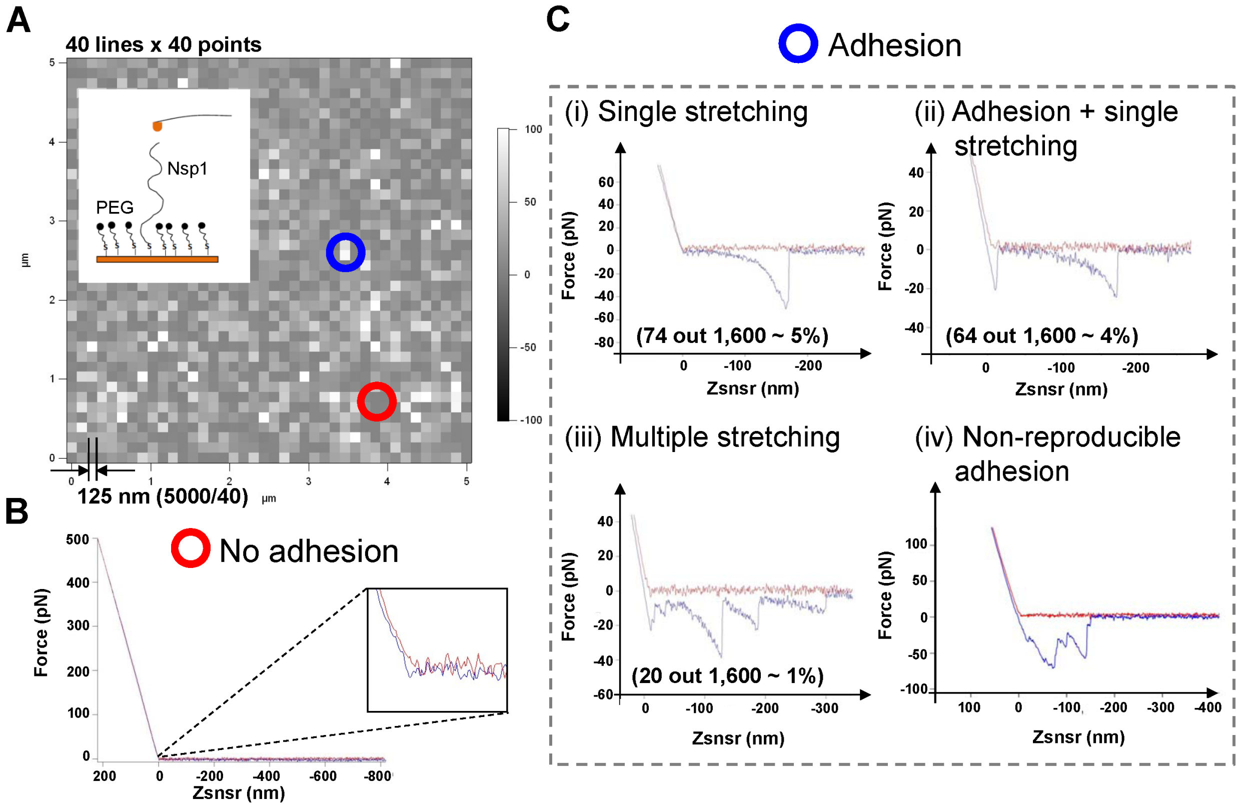

**Figure S7. AFM - Data fitting using the Worm Like Chain model.**

Retraction curves during sNsp1 stretching in PBS are plotted as force, *F(x)*, versus sample-cantilever distance, *x*, and interpolated with the worm-like chain (WLC) model. Examples are reported for (A) single sNsp1 stretching with no initial adhesion peak (multiple runs are shown); (B) sNsp1 stretching with initial adhesion peak; and (C) multiple stretching events. The fitting line from the WLC model is also reported.

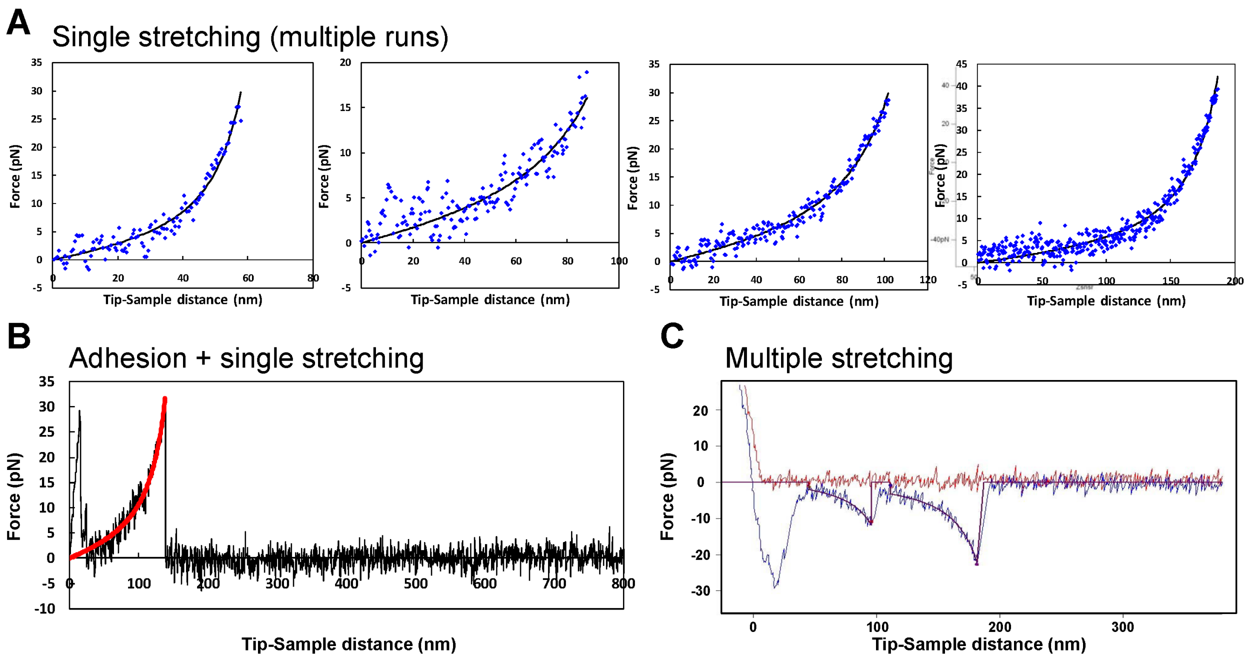

**Figure S8. AFM - Nsp1FG pulling experiment in TBT buffer.**

Plots of the WLC model fitting parameters persistence length, $L_p$ (blue), and contour length, $L_C$ (green), versus the adhesion force, $F_{ad}$, from pulling sNsp1 at 1 μm/s in TBT buffer. Histograms (grey) represent the frequencies of $L_p$, $L_C$ and $F_{ad}$. In the $L_p$ plot, the average (solid line) ± 1 standard deviation (dash line) are also reported.

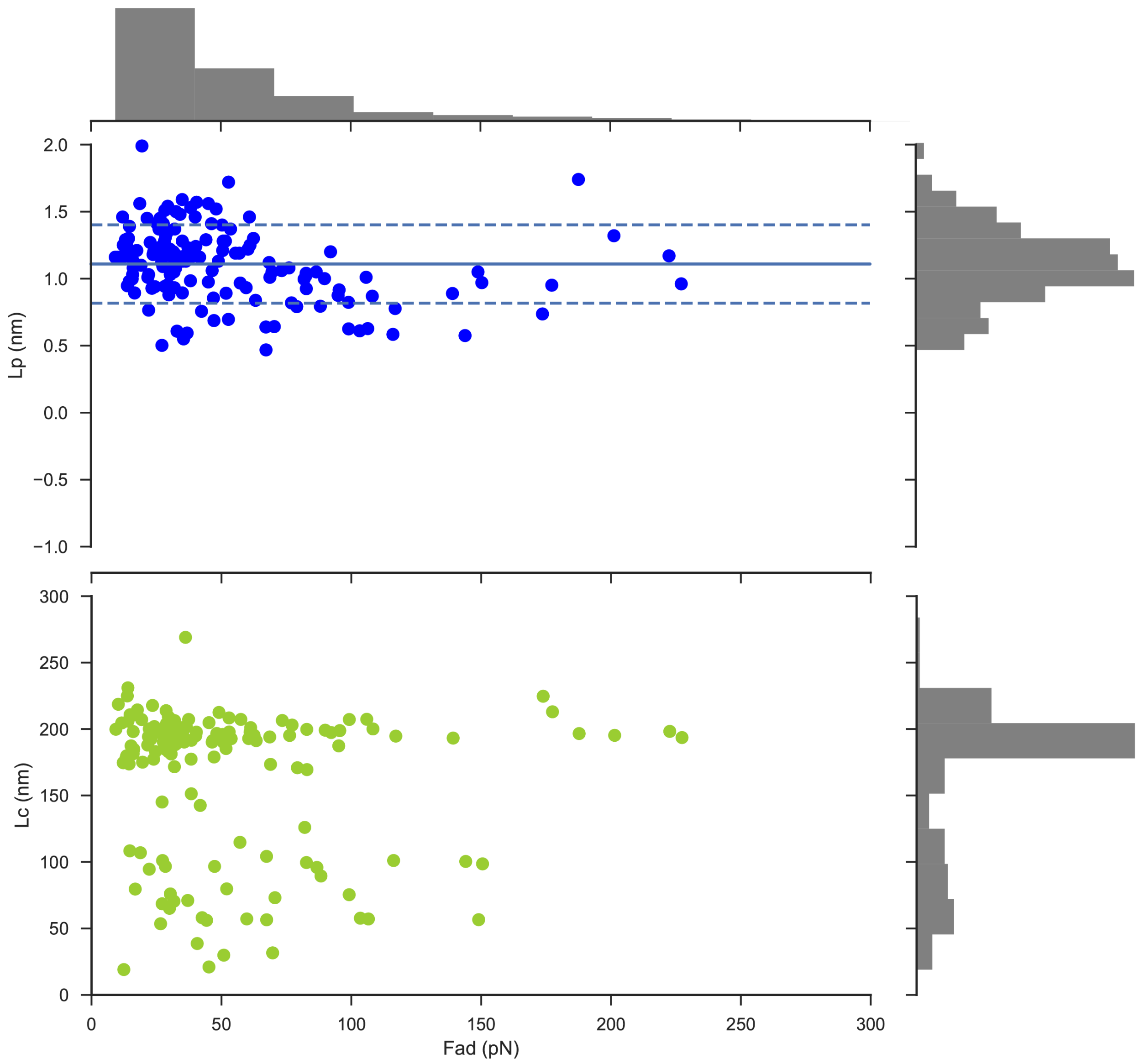

**Figure S9. AFM - Stretching Nsp1 in PBS at different pulling rates.**

Plots of the WLC model fitting parameters persistence length, $L_p$ (blue, top row), and contour length, $L_C$ (green, bottom row), versus the adhesion force, $F_{ad}$, from pulling sNsp1 at (A) 3 μm/s, (B) 2 μm/s and (C) 1 μm/s in PBS buffer. Histograms (grey) represent the frequencies of $L_p$, $L_C$ and $F_{ad}$. In the $L_p$ plots, the average (solid line) ± 1 standard deviation (dash line) are also reported.

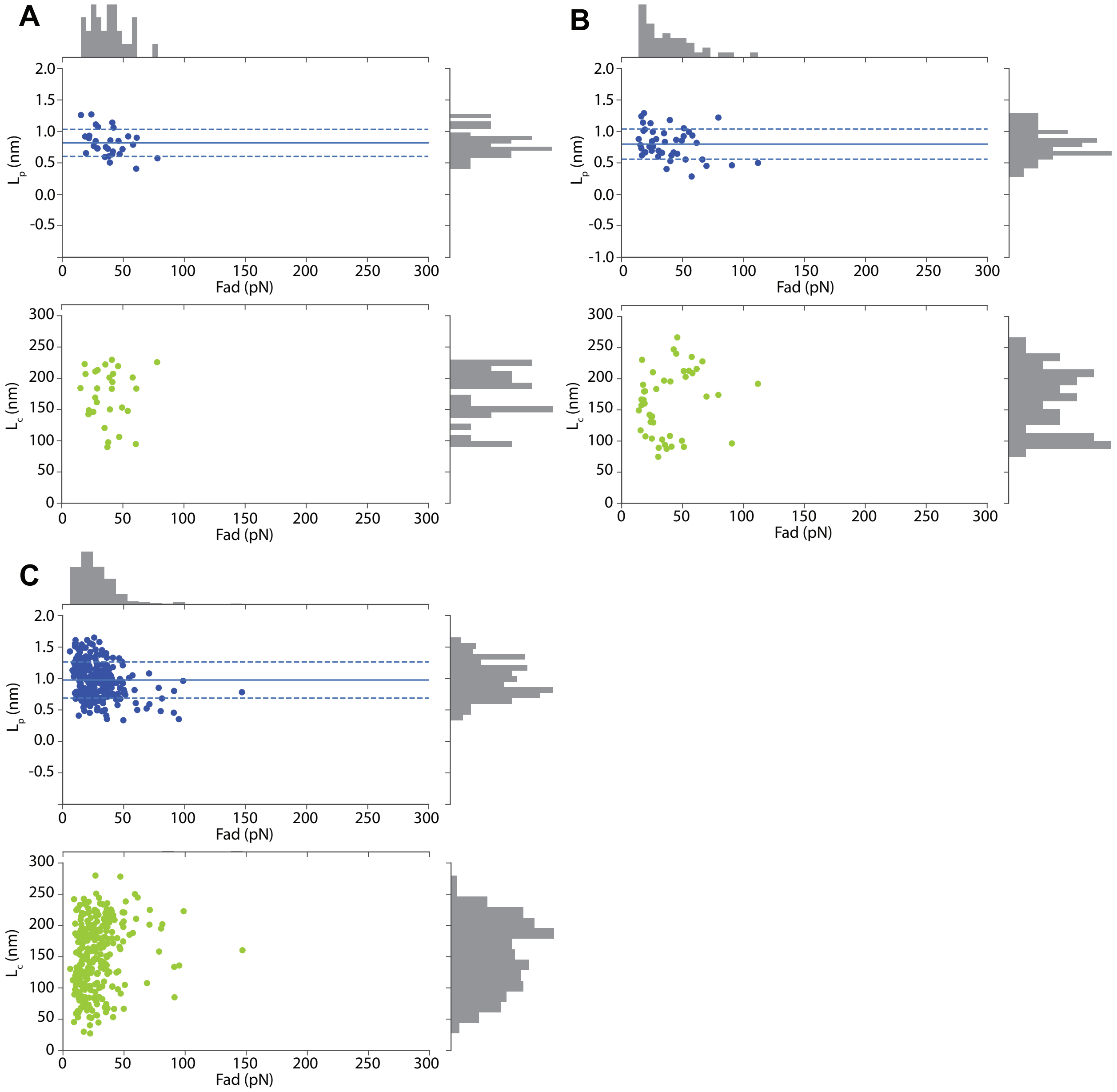

**Figure S10. QCM-D - GST-Kap95 binding with varying binding time.**

The data below show 10 s, 30 s, 1 min, 2 min, and 60 min binding of 1 μM GST-Kap95 on an Nsp1FG-modified sensor, followed by a dissociation phase. The curvature of the binding curves indicates that equilibrium was never reached even after 60 min of binding time. $\Delta F$ and $\Delta D$ indicate the change in resonance frequency and dissipation for the 5th overtone, respectively.

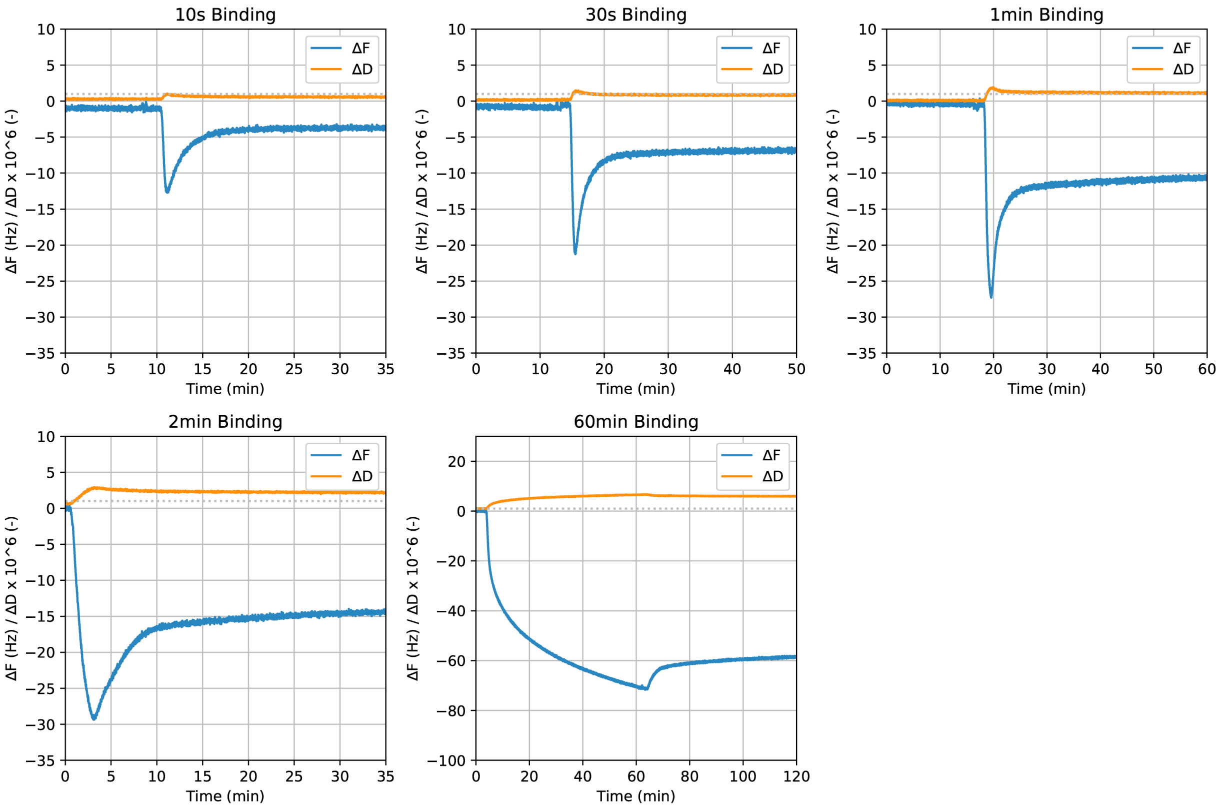

**Figure S11. SPR - Non-specific binding on a bare surface passivated with beta-mercaptoethanol.**

Kap95 bindings on $FSFG_6$ (A-D) and a bare SPR surface passivated with beta-mercaptoethanol (E-H) are shown. Different rows indicate different lengths of the association phase: (A, E) 15 s, (B, F) 30 s, (C, G) 60 s, and (D, H) 120 s. While the SPR response on $FSFG_6$ surface was dependent on Kap95 concentration, it was not for the unconjugated, plain surface. The response came down to zero upon the onset of dissociation phase, suggesting that the plain surface was inert to Kap95.

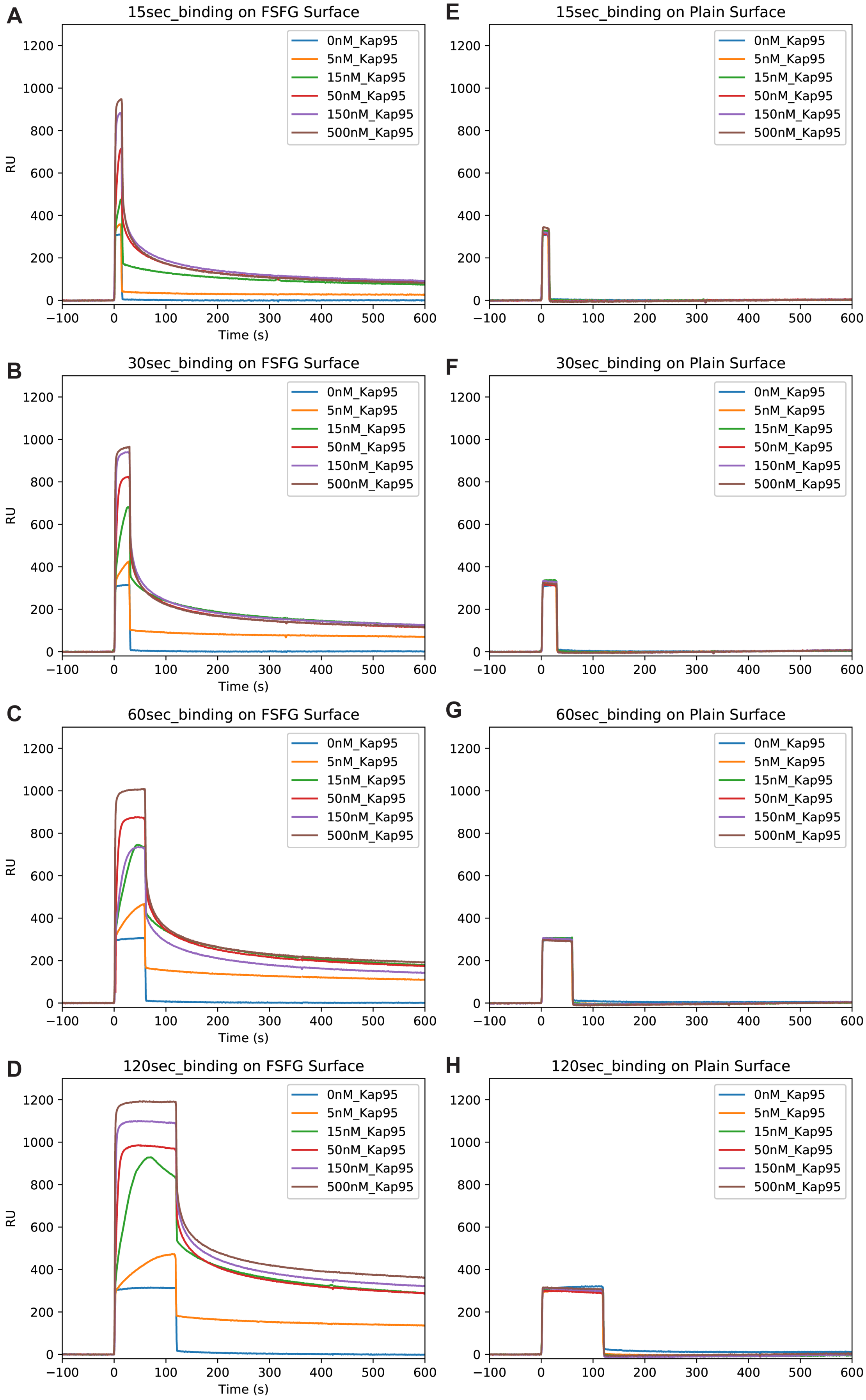
A
15sec_binding on FSFG Surface
E
15sec_binding on Plain Surface
B
30sec_binding on FSFG Surface
F
30sec_binding on Plain Surface
C
60sec_binding on FSFG Surface
G
60sec_binding on Plain Surface
D
120sec_binding on FSFG Surface
H
120sec_binding on Plain Surface
0nM_Kap95
5nM_Kap95
15nM_Kap95
50nM_Kap95
150nM_Kap95
500nM_Kap95
RU
Time (s)

**Figure S12. SPR - Kap95 binding experiments.**

Both association and dissociation phases of the Kap95 binding experiments are shown below. Rows indicate different $FSFG_6$ densities on the surface, and the columns indicate lengths of the association phase. The analysis of the dissociation phase from these experiments is shown in Fig. 4B.

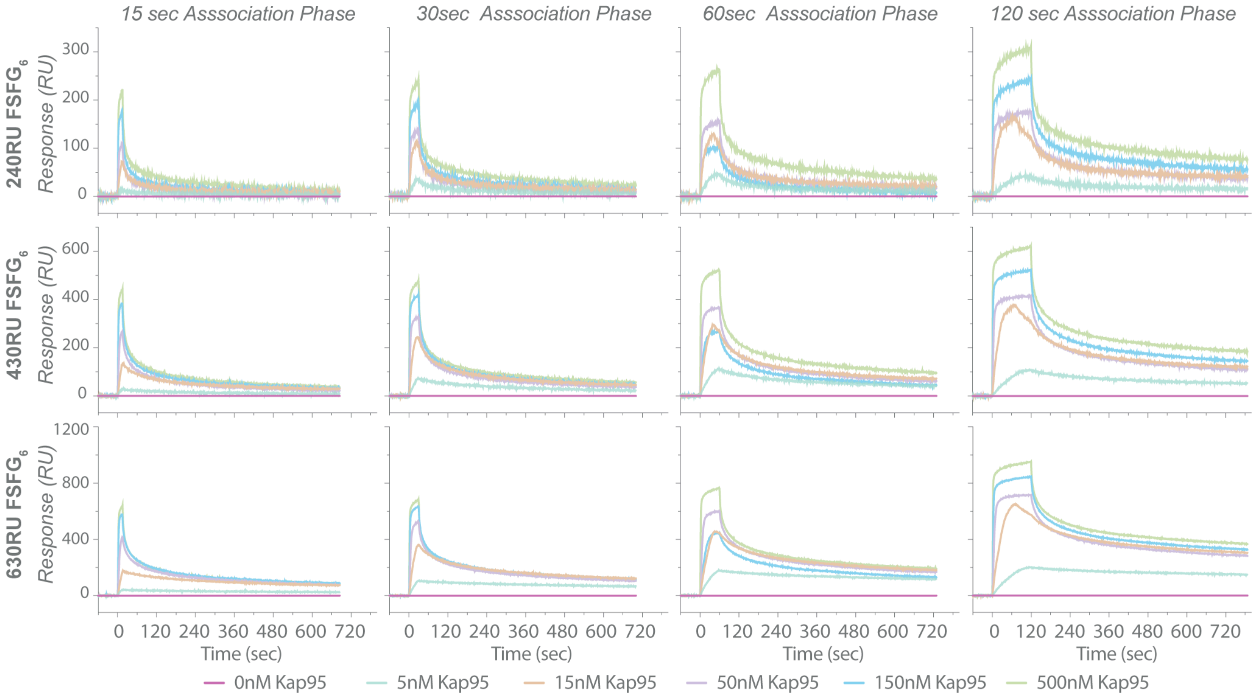

**Figure S13. SPR - Numerical simulation of SPR binding curves.**

Build-up of analyte concentration in an SPR chamber was simulated under different conditions (S1 Text and S1 Table). (A-B) Comparison of analyte build-up in the presence and absence of surface ligand – i.e. (+) and (-) adsorption (ads). (A) It takes ~2-3 s to fully replace the SPR chamber volume. (B) Expansion of the first 0.5 s of (A) shows an additional mixing lag in the presence of surface ligand relative to the bare surface. Comparisons of (C-D) analyte build-up and (E-F) bound analyte at different flow rates. Even at flow rates that are an order of magnitude faster than those used in our SPR experiments (i.e., 100 μl/min) and with smaller model proteins that diffuse faster than those used in our experiments, the simulations indicate mixing lags approaching a few tenths of a second. This time scale is orders of magnitude larger than the time scale of FG-Kap95 interactions, suggesting that mass transport limitation was inevitable with our SPR setting. The reported values are averaged concentration values at the outlet of the SPR chamber for (A-D) and the total amount of analyte bound over the lower SPR reaction surface for (E-F) (see model description). (B), (D) and (F) are magnifications of (A), (C) and (E), respectively.

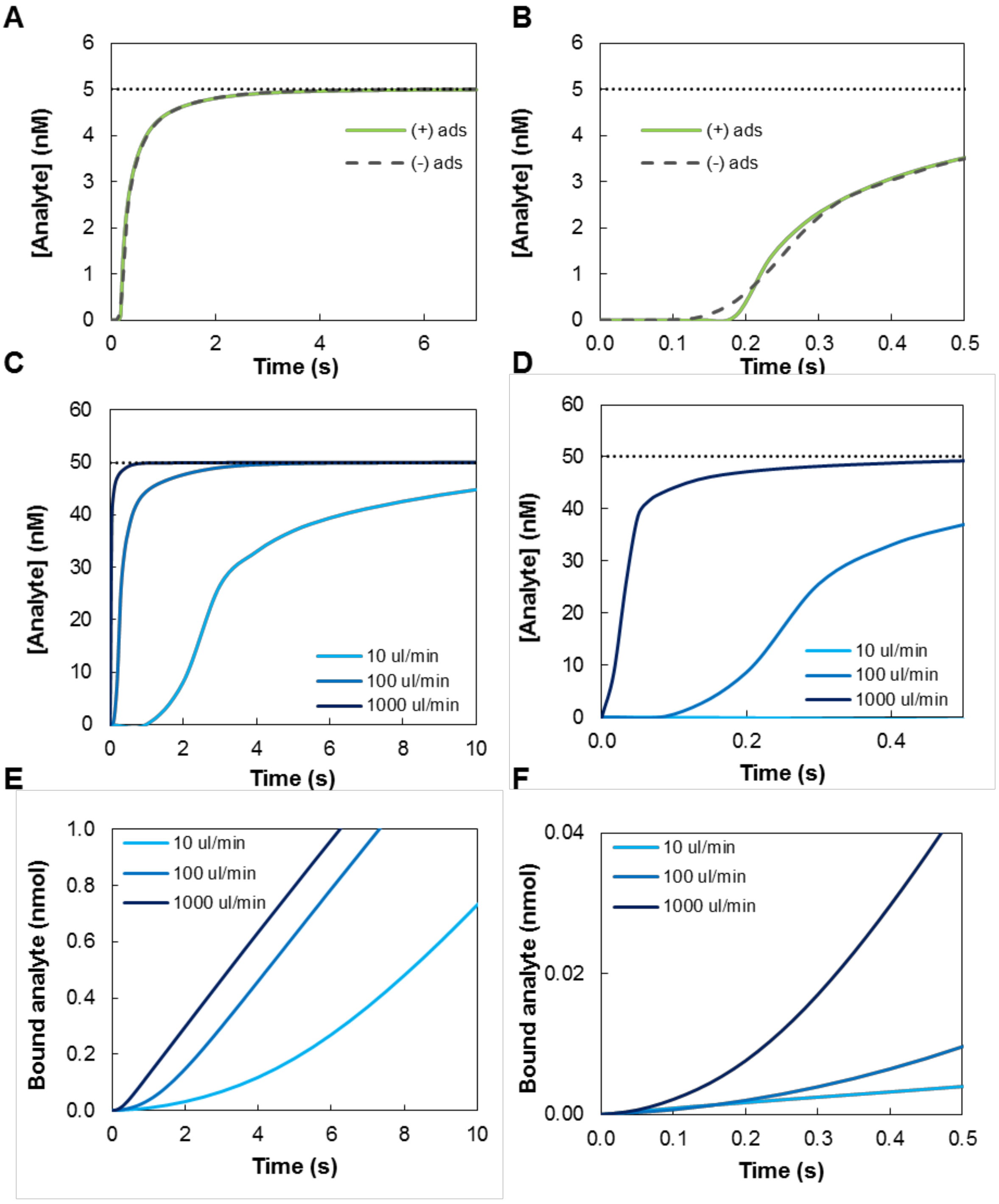
A
[Analyte] (nM)
Time (s)
(+) ads
(-) ads
B
[Analyte] (nM)
Time (s)
(+) ads
(-) ads
C
[Analyte] (nM)
Time (s)
10 ul/min
100 ul/min
1000 ul/min
D
[Analyte] (nM)
Time (s)
10 ul/min
100 ul/min
1000 ul/min
E
Bound analyte (nmol)
Time (s)
10 ul/min
100 ul/min
1000 ul/min
F
Bound analyte (nmol)
Time (s)
10 ul/min
100 ul/min
1000 ul/min

**Figure S14. SPR - Fitting various kinetic models to binding curves.**

SPR binding curves were fit to various kinetic models: one-to-one (A), one-to-one with MTL (B), two state models (C), heterogeneous ligand (D), and heterogeneous analyte (E) Binding curves on surfaces with different $FSFG_6$ densities are shown (columns (i) 425 RU, (ii) 550 RU, (iii) 791 RU). Data here are from a replicate experiment of the one shown in Fig. 4. Solid and dotted lines indicate the obtained sensorgrams and the respective fitted curves, respectively Kap95 concentrations are indicated in different colors. See S2 Table for the fitted parameters values.

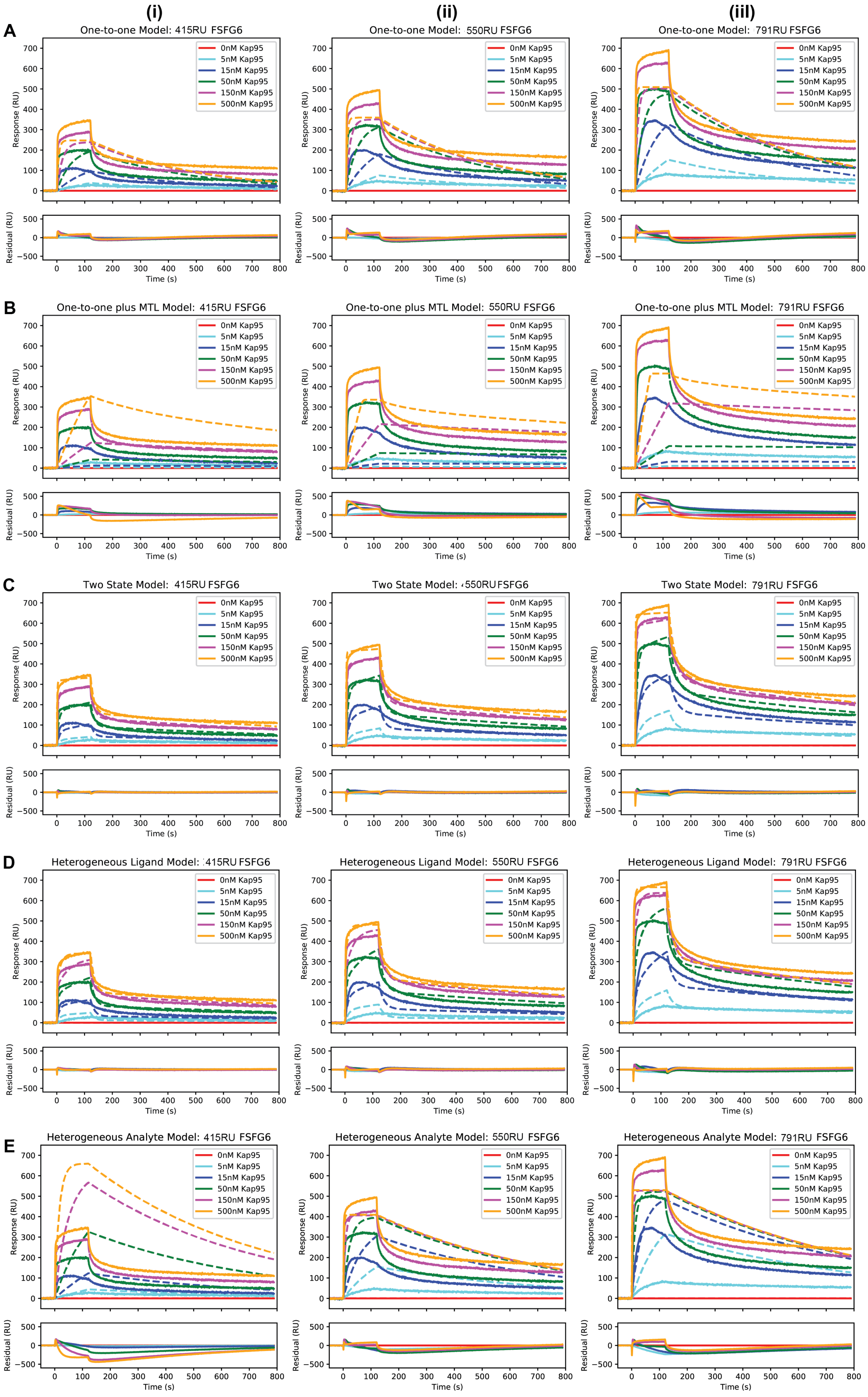

(i)
(ii)
(iii)
A
One-to-one Model: 415RU FSFG6
One-to-one Model: 550RU FSFG6
One-to-one Model: 791RU FSFG6
B
One-to-one plus MTL Model: 415RU FSFG6
One-to-one plus MTL Model: 550RU FSFG6
One-to-one plus MTL Model: 791RU FSFG6
C
Two State Model: 415RU FSFG6
Two State Model: 550RU FSFG6
Two State Model: 791RU FSFG6
D
Heterogeneous Ligand Model: 415RU FSFG6
Heterogeneous Ligand Model: 550RU FSFG6
Heterogeneous Ligand Model: 791RU FSFG6
E
Heterogeneous Analyte Model: 415RU FSFG6
Heterogeneous Analyte Model: 550RU FSFG6
Heterogeneous Analyte Model: 791RU FSFG6
0nM Kap95
5nM Kap95
15nM Kap95
50nM Kap95
150nM Kap95
500nM Kap95
Response (RU)
Residual (RU)
Time (s)

**Figure S15. QCM-D - Conjugation of $FSFG_6$ and $SSSG_6$ to silica sensors.**

The changes in frequency ($\Delta F$) and energy dissipation ($\Delta D$) for several overtones were recorded in real time during (A) $FSFG_6$ and (B) $SSSG_6$ conjugations. The overlapping of the different harmonics is indicative of the rigid nature of these layers. (C-D) Sauerbrey model was used to calculate the layer thickness, for different layer densities (1,100, 1,200 and 1,300 kg/m$^3$).

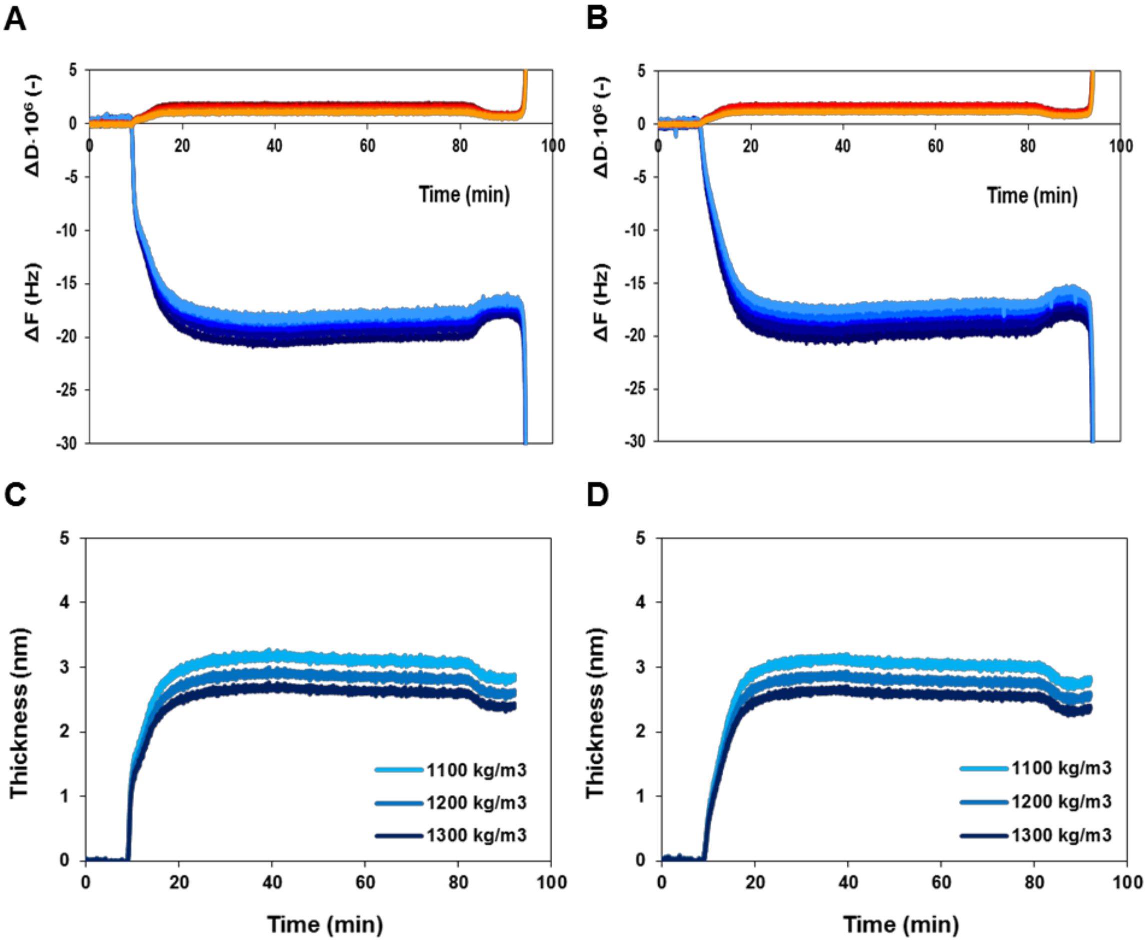

**Table S1. SPR - Numerical simulation parameters.**

Summary of the parameters used for the SPR numerical simulations in S13 Fig.

| Parameter description | Value | Units |
|---|---|---|
| ***SPR geometry*** | | |
| Channel depth, D | $1{\cdot}10^{-3}$ | m |
| Channel width, W | $1{\cdot}10^{-3}$ | m |
| Channel height, H | $5{\cdot}10^{-4}$ | m |
| Chamber volume, D·W·H | $5{\cdot}10^{-10}$ | $m^3$ |
| Surface area for adsorption, D·W | $1{\cdot}10^{-6}$ | $m^2$ |
| Outlet surface area, D·H | $5{\cdot}10^{-7}$ | $m^2$ |
| ***Flow properties*** | | |
| Inlet velocity[a] | $(0.3 – 3 – 30){\cdot}10^{-3}$ | m/s |
| $H_2O$ concentration | 55,000 | $mol/m^3$ |
| $H_2O$ viscosity | $8.9{\cdot}10^{-4}$ | Pa·s |
| $H_2O$ molecular weight | 18 | g/mol |
| $H_2O$ density | 1,000 | $kg/m^3$ |
| Temperature | 298 | K |
| ***Ligand*** | | |
| Molecular weight, $MW_L$ | 28,771 | g/mol |
| Experimental RU[b] | 500 - 1,500 | - |
| Total binding sites[c] | $(RU/1,000)/MW_L$ | $mol/m^2$ |
| ***Analyte*** | | |
| Molecular weight, $MW_A$ | 15,565 | g/mol |
| Inlet concentration | $5{\cdot}10^{-5}$ | $mol/m^3$ |
| ***Binding*** | | |
| Adsorption rate constant | $2{\cdot}10^{2}$ | $mol/(m^3{\cdot}s)$ |
| Desorption rate constant | $2{\cdot}10^{-3}$ | 1/s |

[a] The feed velocity was 3 mm/s, corresponding to the experimental 100 µL/min, for S13A-B Fig, while it was varied from 0.3 to 30 mm/s for S13C-F Fig.

[b] Experimental SPR response unit (RU) signal was 1,500 (i.e. high ligand density) for S13A-B Fig and 500 (i.e. medium ligand density) for S13C-F Fig.

[c] It was calculated using the correlation 1 RU = 1,000 $ng/mm^2$.

**Table S2. SPR - Fitting kinetic and equilibrium models to binding curves**

SPR binding curves (solid lines in S14 Fig) were fit to various kinetic models: one-to-one, one-to-one with MTL, heterogeneous ligands, heterogeneous analytes, and two state models. Langmuir isotherm analysis was also included. FG density refers to the amount of $FSFG_6$ conjugated on the SPR sensors: low (415 RU), medium (550 RU) and high (791 RU). Binding time indicate the length of the association phase: 15, 30, 60 and 120 seconds.

*One-to-one model*

| FG Density | Binding Time (s) | ka ($10^5$ 1/Ms) | kd ($10^{-3}$ 1/s) | KD ($10^{-9}$ M) | $R_{max}$ (RU) | $Chi^2$ (RU) |
|---|---|---|---|---|---|---|
| Low | 15 | 10.80 | 3.70 | 3.42 | 142.4 | 516 |
| | 30 | 5.32 | 3.01 | 5.65 | 182.6 | 711 |
| | 60 | 5.49 | 3.01 | 5.49 | 218.0 | 1173 |
| | 120 | 3.21 | 2.61 | 8.14 | 251.5 | 1154 |
| Medium | 15 | 20.70 | 3.03 | 1.46 | 222.4 | 974 |
| | 30 | 8.10 | 2.65 | 3.27 | 279.8 | 1403 |
| | 60 | 8.73 | 2.75 | 3.15 | 319.5 | 2349 |
| | 120 | 4.59 | 2.50 | 5.45 | 362.8 | 2383 |
| High | 15 | 24.10 | 2.09 | 0.87 | 386.7 | 1643 |
| | 30 | 9.97 | 1.99 | 1.99 | 442.5 | 2364 |
| | 60 | 12.9 | 2.20 | 1.70 | 470.7 | 4077 |
| | 120 | 6.82 | 2.19 | 3.20 | 512.5 | 4249 |

*One-to-one plus mass transport limitation model*

| FG Density | Binding Time (s) | ka ($10^5$ 1/Ms) | kd ($10^{-3}$ 1/s) | KD ($10^{-9}$ M) | $R_{max}$ (RU) | kt ($10^4$ RU/Ms) | $Chi^2$ (RU) |
|---|---|---|---|---|---|---|---|
| Low | 15 | 9.75 | 6.53 | 6.70 | 208.0 | 1 | 2129 |
| | 30 | 42.90 | 4.49 | 1.05 | 74.6 | 991 | 1642 |
| | 60 | 9.05 | 4.09 | 4.52 | 227.6 | 877 | 2973 |
| | 120 | 1.15 | 3.26 | 28.30 | 434.2 | 849 | 5035 |
| Medium | 15 | 25.60 | 4.65 | 1.82 | 166.8 | 1 | 6896 |
| | 30 | 44.50 | 3.30 | 0.74 | 99.0 | 1560 | 5199 |
| | 60 | 15.50 | 3.86 | 2.49 | 271.2 | 1210 | 7340 |
| | 120 | 8.51 | 3.18 | 3.74 | 338.2 | 1290 | 7949 |
| High | 15 | 3.58 | 3.35 | 9.36 | 754.9 | 1 | 27110 |
| | 30 | 35.70 | 2.70 | 0.76 | 127.6 | 3850 | 17795 |
| | 60 | 28.30 | 3.14 | 1.11 | 315.9 | 1840 | 21187 |
| | 120 | 16.60 | 2.75 | 1.65 | 466.0 | 1830 | 20928 |

*Heterogeneous analytes model*

| FG Density | Binding Time (s) | ka ($10^5$1/Ms) | kd ($10^{-3}$ 1/s) | KD ($10^{-9}$ M) | $R_{max}$ (RU) | $ka_2$ ($10^5$1/Ms) | $kd_2$ ($10^{-3}$ 1/s) | $KD_2$ ($10^{-9}$ M) | $Chi^2$ (RU) |
|---|---|---|---|---|---|---|---|---|---|
| Low | 15 | 1.67 | 3.27 | 19.60 | 515.0 | 1.67 | 3.27 | 19.60 | 6092 |
| | 30 | 21.9 | 2.24 | 1.03 | 80.5 | 21.90 | 2.24 | 1.03 | 1341 |
| | 60 | 4.66 | 2.04 | 4.38 | 282.1 | 4.66 | 2.04 | 4.38 | 3784 |
| | 120 | 0.612 | 1.63 | 26.60 | 677.2 | 0.61 | 1.63 | 26.60 | 27094 |
| Medium | 15 | 12.3 | 2.33 | 1.89 | 258.3 | 12.30 | 2.33 | 1.89 | 1710 |
| | 30 | 22.5 | 1.65 | 0.73 | 107.4 | 22.50 | 1.65 | 0.73 | 3789 |
| | 60 | 7.89 | 1.93 | 2.45 | 363.5 | 7.89 | 1.93 | 2.45 | 5473 |
| | 120 | 4.36 | 1.59 | 3.65 | 411.3 | 4.36 | 1.59 | 3.65 | 7967 |
| High | 15 | 4.63 | 1.68 | 3.62 | 641.9 | 4.63 | 1.68 | 3.62 | 10615 |
| | 30 | 18.8 | 1.35 | 0.72 | 157.8 | 18.80 | 1.35 | 0.72 | 12260 |
| | 60 | 14.9 | 1.57 | 1.05 | 380.1 | 14.90 | 1.57 | 1.05 | 5768 |
| | 120 | 8.4 | 1.37 | 1.64 | 530.4 | 8.40 | 1.37 | 1.64 | 13870 |

*Heterogeneous ligands model*

| FG Density | Binding Time (s) | ka ($10^5$1/Ms) | kd ($10^{-3}$ 1/s) | KD ($10^{-9}$ M) | $R_{max}$ (RU) | $ka_2$ ($10^5$1/Ms) | $kd_2$ ($10^{-3}$ 1/s) | $KD_2$ ($10^{-9}$ M) | $Chi^2$ (RU) |
|---|---|---|---|---|---|---|---|---|---|
| Low | 15 | 31.90 | 85.4 | 26.70 | 243.0 | 4.27 | 1.19 | 2.77 | 83 |
| | 30 | 20.10 | 80.2 | 39.90 | 251.9 | 2.61 | 1.00 | 3.84 | 87 |
| | 60 | 38.90 | 82.7 | 21.30 | 228.7 | 1.95 | 0.87 | 4.43 | 159 |
| | 120 | 1.38 | 0.7 | 5.12 | 152.9 | 33.20 | 71.90 | 21.70 | 125 |
| Medium | 15 | 20.10 | 1.5 | 0.76 | 142.6 | 20.50 | 74.90 | 36.50 | 222 |
| | 30 | 5.72 | 1.1 | 1.99 | 183.2 | 18.30 | 68.20 | 37.30 | 248 |
| | 60 | 4.48 | 1.0 | 2.14 | 198.5 | 39.50 | 68.80 | 17.40 | 467 |
| | 120 | 2.17 | 0.7 | 3.32 | 219.6 | 35.70 | 57.70 | 16.20 | 368 |
| High | 15 | 15.10 | 54.6 | 36.20 | 452.5 | 25.60 | 1.28 | 0.50 | 360 |
| | 30 | 9.39 | 1.0 | 1.09 | 317.6 | 14.50 | 57.00 | 39.30 | 373 |
| | 60 | 11.60 | 1.0 | 0.83 | 310.4 | 26.10 | 60.00 | 23.00 | 918 |
| | 120 | 4.86 | 0.8 | 1.60 | 321.8 | 31.90 | 54.10 | 17.00 | 823 |

*Two state model*

| FG Density | Binding Time (s) | ka ($10^5$1/Ms) | kd ($10^{-3}$ 1/s) | KD ($10^{-9}$ M) | $R_{max}$ (RU) | $ka_2$ ($10^{-3}$ 1/s) | $kd_2$ ($10^{-3}$ 1/s) | $Chi^2$ (RU) |
|---|---|---|---|---|---|---|---|---|
| Low | 15 | 18.60 | 102.0 | 54.7 | 357.6 | 12.10 | 1.60 | 82 |
| | 30 | 11.80 | 85.5 | 72.4 | 384.4 | 9.91 | 1.25 | 75 |
| | 60 | 18.40 | 95.4 | 51.8 | 372.0 | 7.17 | 1.05 | 127 |
| | 120 | 14.70 | 78.6 | 53.3 | 352.7 | 5.24 | 0.86 | 78 |
| Medium | 15 | 28.20 | 55.7 | 19.8 | 453.6 | 12.80 | 1.80 | 194 |
| | 30 | 14.30 | 56.9 | 39.8 | 521.7 | 10.60 | 1.35 | 177 |
| | 60 | 22.20 | 64.0 | 28.8 | 508.4 | 7.42 | 1.09 | 316 |
| | 120 | 17.00 | 60.1 | 35.3 | 492.5 | 5.29 | 0.87 | 208 |
| High | 15 | 29.90 | 28.7 | 9.6 | 639.8 | 13.50 | 1.79 | 428 |
| | 30 | 15.40 | 33.4 | 21.7 | 716.3 | 11.20 | 1.28 | 369 |
| | 60 | 25.20 | 36.0 | 14.3 | 683.6 | 7.65 | 1.05 | 801 |
| | 120 | 19.30 | 39.5 | 20.5 | 667.4 | 5.40 | 0.84 | 523 |

*Langmuir isotherm*

| FG Density | Binding Time (s) | KD ($10^{-9}$ M) | $R_{max}$ (RU) | $Chi^2$ (RU) |
|---|---|---|---|---|
| Low | 15 | 64.2 | 382.9 | 218 |
| | 30 | 76.8 | 397.9 | 345 |
| | 60 | 43.3 | 375.5 | 326 |
| | 120 | 41.9 | 357.0 | 109 |
| Medium | 15 | 59.3 | 580.4 | 1293 |
| | 30 | 63.5 | 585.4 | 1382 |
| | 60 | 30.8 | 537.1 | 1042 |
| | 120 | 30.4 | 505.7 | 437 |
| High | 15 | 64.5 | 892.1 | 2203 |
| | 30 | 54.7 | 860.3 | 4405 |
| | 60 | 22.6 | 764.5 | 3062 |
| | 120 | 22.4 | 708.9 | 1519 |
